# Structure-guided discovery and engineering of miniature CRISPR-Cas12m for epigenome editing

**DOI:** 10.64898/2026.03.26.714355

**Authors:** Tao Yu, Meng Ji, Donglin Yu, Zhao Guan, Rongyi Zhu, Yunpeng Jiang, Zhiyi Yang, Lizhen Qiu, Jiawei Mu, Fengbiao Mao, Kuanhui Xiang, Lin Bai, Kailong Li

## Abstract

CRISPR-based epigenome editing represents a programmable strategy to precisely modulate gene expression, holding immense promise for therapeutic applications. However, the large size of the dCas proteins substantially impedes the delivery via adeno-associated virus (AAV) vectors. Here, through iterative bioinformatics analysis, structure-guided predictions, and functional assays, we identified and characterized PmCas12m, a novel miniature subtype V-M CRISPR-Cas12m. PmCas12m exhibited flexible 5’-YTN-3’ PAM-dependent recognition and robust double-stranded DNA binding properties, while lacking DNA cleavage activity, thus positioning it as an ideal tool for epigenome editing. Cryogenic electron microscopy (cryo-EM) structures of PmCas12m unveiled its unique molecular mechanism of DNA binding facilitating interference. Guided by these structural insights, we employed deep mutational scanning (DMS) and protein engineering to develop xCas12m, a hypercompact variant with highly potent and specific epigenome editing capabilities in human cells. We further constructed the xCas12m-CRISPRoff platform in a single AAV vector, which achieved durable epigenetic silencing and effective inhibition of hepatitis B virus (HBV) infection in a mouse model. Collectively, these findings establish xCas12m as a versatile epigenome editing platform with transformative potential for treating diseases, paving the way for clinical translation of epigenetic therapies.

## Introduction

Epigenetic regulatory mechanisms play a central role in nearly all cellular processes, while their dysregulation manifests in aberrant gene expression and diseases like viral infection^1^. The CRISPR-based epigenome editing system offers a versatile array of epigenetic effector domains that can be fused to nuclease-dead Cas protein (dCas), generating a rich toolset for transcriptional regulation (CRISPRa and CRISPRi)^2,3^. This technology combines the precision of CRISPR with epigenetic mark rewriting, offering a tunable and reversible gene regulation approach without altering the DNA sequences, thus holding immense potential for therapeutic applications^2–5^. However, the large size of the commonly used dCas proteins poses challenges for their efficient packing in cargo-size-limited *in vivo* delivery vehicles like AAV, and the fusion of additional domains further increases the molecular weight, impeding clinical translation^6–8^. As such, tremendous efforts have been made to overcome this obstacle. One approach involves splitting the reagents into two AAV vectors, but this lowers delivery efficiency as both vectors must transduce the same cell^6,7^. An alternative strategy focuses on engineering miniature dCas proteins, though challenges remain in ensuring their targeting specificity and functional activity^6,8^. As such, the development of more compact and efficient dCas proteins that can be packaged in a single AAV is crucial to advance the feasibility and therapeutic potential of epigenome editing.

Recently, researchers have discovered three novel type V CRISPR–Cas systems (Cas12c^9^, Cas12k^10,11^, and Cas12m^12,13^), as well as a family of TnpB-like nuclease-dead repressors (TldRs)^14^. These systems are characterized by their lack of DNA cleavage activity while retaining RNA-guided DNA binding capabilities. The identification of naturally occurring nuclease-deficient CRISPR-Cas proteins raises the possibility of their unique potential as valuable tools for epigenome editing. By functioning inherently as efficient DNA-targeting platforms, these proteins may offer a streamlined and advantageous alternative for achieving precise epigenetic regulation. However, most of these proteins present unique challenges that restrict their application potential. For instance, the relatively large size of Cas12c complicates its use in AAV-mediated gene delivery^9^. Cas12k exhibits minimal to no activity in eukaryotic cells^11^, and the activity of TldRs in human cells remains ambiguous^14^. In contrast, Cas12m stably binds to the target dsDNA in human cells, together with the small size and abundant putative homologs in prokaryotes^13^, it represents an underexplored enormous source of miniature genome editors.

Hepatitis B virus (HBV) infection remains a major global health problem, with chronic infection affecting millions of individuals worldwide^15,16^. Current antiviral therapies, particularly nucleos(t)ide analogues (NAs), are effective at suppressing HBV replication and reducing viral load. However, these therapies do not provide a complete cure and typically require lifelong treatment^17,18^. CRISPR/Cas9 shows promise for editing the HBV genome to disrupt infection and replication but is hindered by unintended DNA breakage ^19–24^. In contrast, epigenetic editing has emerged as a safer alternative for antiviral therapies, offering precise control of gene expression without inducing DNA breakage, thereby reducing the risk of genomic damage. More importantly, increasing evidence shows that epigenetic modification of the viral genome is essential for the regulation of viral activity, positioning the HBV epigenome as an attractive therapeutic target^25–27^.

In this study, we developed and applied a structure-guided search pipeline to identify, characterize and engineer a miniature CRISPR-Cas12m protein (xCas12m). xCas12m exhibited highly efficient epigenome editing capabilities with minimal off-target effects in human cells. We further constructed the xCas12m-CRISPRoff platform, which achieved durable epigenetic silencing and effective inhibition of HBV activity in a mouse model with a single AAV vector. Together, our work establishes xCas12m as a hypercompact and durable epigenome editing tool with broad therapeutic potential, highlighting its promise as a clinically translatable platform for combating viral infections through targeted epigenetic silencing.

## Results

### Identification and characterization of novel Cas12m proteins via a structure-guided pipeline

To investigate the diversity of Cas12m and discover novel candidates for expanding the CRISPR biology, we developed an automated structure-guided search pipeline to identify nuclease-dead Cas12m proteins harboring inactivating mutations in the RuvC DED motif (Fig. 1a). Utilizing MmCas12m(WP_061006603.1)^13^ as an initial query, we performed a PSI-BLAST search against the NCBI non-redundant database and then constructed multiple HMM profiles^28^ of the corresponding representative protein domain architecture based on the sequence alignments. This search strategy, cooperated with additional CRISPR array analysis^29^, identified 30 Cas12m candidates with inactive RuvC nuclease activity from various bacterial families that potentially harbor active CRISPR-Cas systems in the NCBI RefSeq database. Given that a protein’s biological function is closely linked to its structural properties, we hypothesized that proteins with comparable structures are likely to exhibit similar functions. Therefore, we employed AlphaFold 3^30^ to predict the 3D structures of the CRISPR-Cas12m candidates and compared these predicted structural architectures with that of MmCas12m using the DALI web server^31^. Ultimately, we selected the top 15 candidates for further functional testing (Supplementary Table 1).

**Fig. 1.**
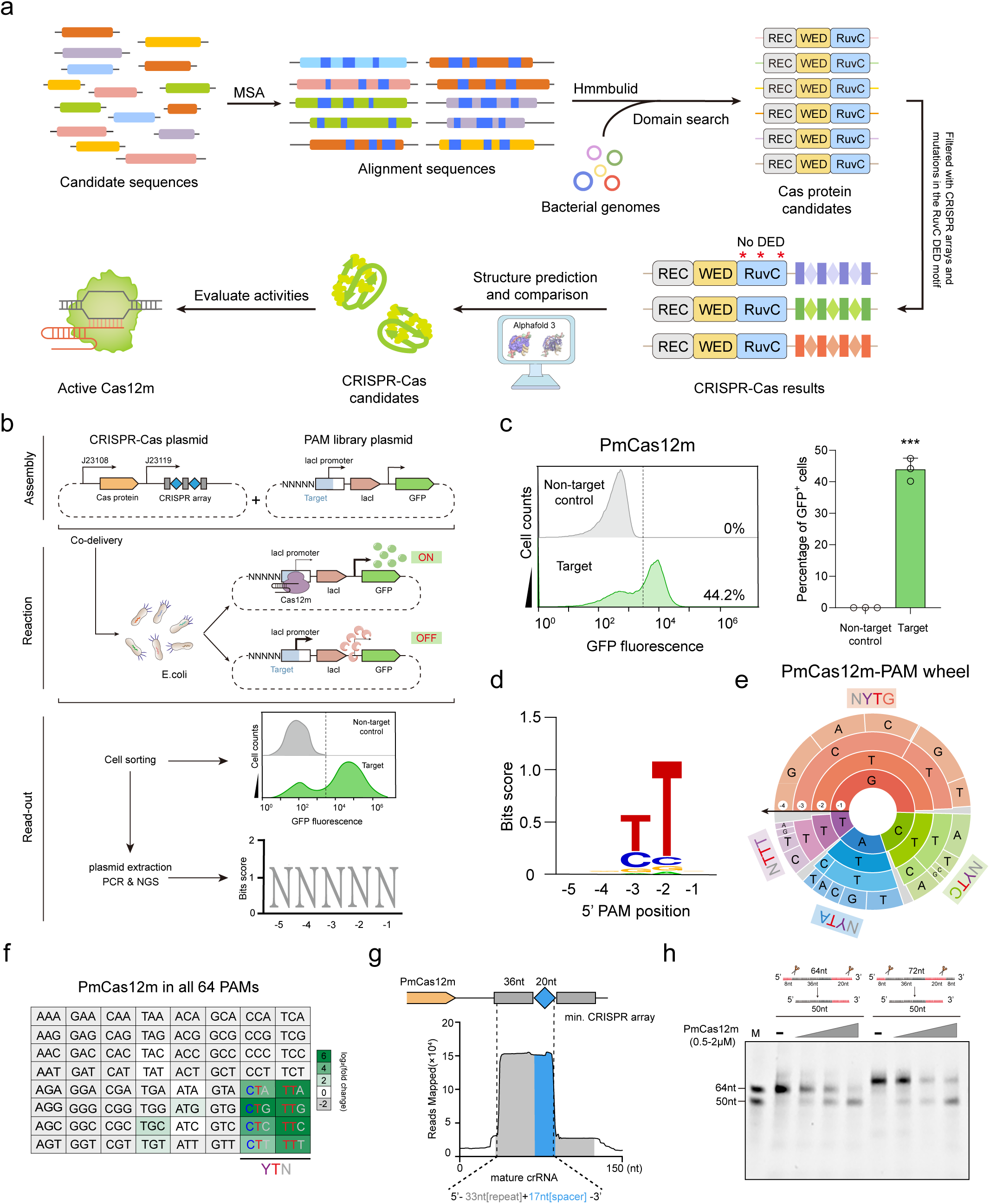
Discovery and characterization of Cas12m homologs. **a,** Schematic illustrating the pipeline for de novo identification of naturally occurring nuclease-dead CRISPR-Cas12m homologs. **b**, *In vivo* PAM screening platform implemented through PAM-SCANR. **c**, Left: Cells harboring functional PAMs that led to GFP fluorescence were sorted by FACS. Right: FACS analysis showing the percentage of GFP-positive cells for PmCas12m. Data represent the mean ± s.d. of three biological replicates. ***P value < 0.001. **d-f**, PAM recognition profiles of PmCas12m with 5-nt (**d)**, 4-nt (**e)** and 3-nt (**f)** PAMs. **g**, Small RNA-seq results mapped onto the minimal CRISPR-PmCas12m system. **h**, Pre-crRNA processing assay demonstrating PmCas12m-mediated cleavage activity.

To explore the potential activity of these Cas12m candidates, we performed the PAM-SCANR assay for PAM determination as previously described (Fig. 1b)^32^. Specifically, we optimized the electroporation conditions in *E. coli* to deliver an all-in-one plasmid expressing both Cas12m and the corresponding RNA components, along with the PAM-SCANR plasmid library containing a 5-base pair (bp) randomized region upstream of the protospacer. Following cultivation, fluorescent cells were isolated and sorted by fluorescence-activated cell sorting (FACS). Notably, functional Cas12m binding to PAM-SCANR plasmids containing valid PAM sequences would result in GFP expression. We applied this assay to screen all 15 candidates and found that PmCas12m, MkCas12m and PpCas12m induced robust reporter activity, with PmCas12m being the most effective one (Fig. 1c and Extended Data Fig. 1a). Deep sequencing of the GFP^+^ plasmids revealed that PmCas12m, MkCas12m and PpCas12m recognized PAM of 5’-YTN-3’ (Y=C or T), 5’-YTTS-3’ (Y=C or T, S=G or C) and 5’-TTV-3’ (V=not T), respectively (Fig. 1d-f and Extended Data Fig. 1b). Given that PmCas12m exhibited the highest binding activity and demonstrated a flexible PAM recognition feature, we selected it for further functional investigation.

To elucidate the essential RNA components of PmCas12m, we conducted small RNA-seq using *E. coli* expressing a minimal CRISPR-Cas12m system. As shown in Fig. 1g, we identified a ∼50-nucleotide (nt) mature crRNA containing a ∼33-nt direct repeat followed by a ∼17 nt guide sequence. Additionally, the *in vitro* pre-crRNA processing assay was conducted using purified PmCas12m and pre-crRNA with a 5’ extension, with or without a 3’ extension. The results indicated that cleavage predominantly occurred after the 2^nd^-3^rd^ nucleotide of the direct repeat, as well as at the 17^th^-18^th^ position of the guide sequence (Fig. 1h), consistent with the findings from the small RNA-seq analysis (Fig. 1g). To confirm the nuclease-dead feature of Cas12m, we performed the PAM depletion assay and found that all three Cas12m proteins were catalytically inactive *in vivo* (Extended Data Fig. 1c). Consistent with these findings, PmCas12m also failed to exhibit cleavage activity *in vitro*, further confirming its nuclease-dead characteristic (Extended Data Fig. 2). Together, these results demonstrated that PmCas12m bound but did not cleave dsDNA with a flexible 5’-YTN-3’ PAM.

### PmCas12m blocks target gene expression via binding to dsDNA in *E. coli*

CRISPR-Cas systems are generally thought to protect prokaryotes from phage or viral infections through their nuclease activity, which enables them to cleave foreign DNA. However, the nuclease-deficient characteristic of PmCas12m raises the question of whether it can function as an effective defense system by binding to specific target sites, similar to other type V CRISPR-Cas systems^9,12,13^. To test this hypothesis, we initially employed a dual-color fluorescence interference assay. In this assay, we used a pTarget-RFP-GFP operon reporter plasmid (addgene#192279) containing a bi-cistronic operon with two reporter genes, GFP and RFP^13,33^. *E. coli* cells were co-transformed with the pTarget-RFP-GFP operon plasmid and PmCas12m plasmid, along with CRISPR array plasmids targeting different sites within the GFP and RFP coding sequences on both strands (Fig. 2a and Supplementary Table 2). If PmCas12m bound to the target DNA without cleaving it, we expected a reduction in the targeted fluorophore signal without causing cell death. As anticipated, when guided by a GFP- or RFP-targeting sgRNA, PmCas12m effectively silenced gene expression while preserving the cell viability, showing no strand preference (Fig. 2b, c). Building on the observation that PmCas12m provided binding-based transcription inhibition when targeting the marker gene, we further investigated whether PmCas12m exhibited interference activity by binding to essential regions involved in plasmid replication and propagation. To examine this, the pTarget-GFP reporter plasmid was transformed into *E. coli* cells harboring the PmCas12m plasmid and CRISPR array plasmid targeting Kan and GFP coding sequences, the origin of replication (ori) region and non-essential sequence on both strands (Fig. 2d). As expected, when sgRNAs were targeted to the essential regions, such as Kan and ori, they disrupted the plasmid replication or repressed the essential gene expression in *E. coli*, leading to cell death or growth defects (Fig. 2d, e and Supplementary Table 2). Importantly, PmCas12m demonstrated no strand bias when targeted to the noncoding regions (Fig. 2d, e). Collectively, these findings suggest that PmCas12m functions as a defense system by blocking the transcription of essential genes, thus providing adaptive immunity against invading mobile genetic elements (MGE) through its binding activity.

**Fig. 2.**
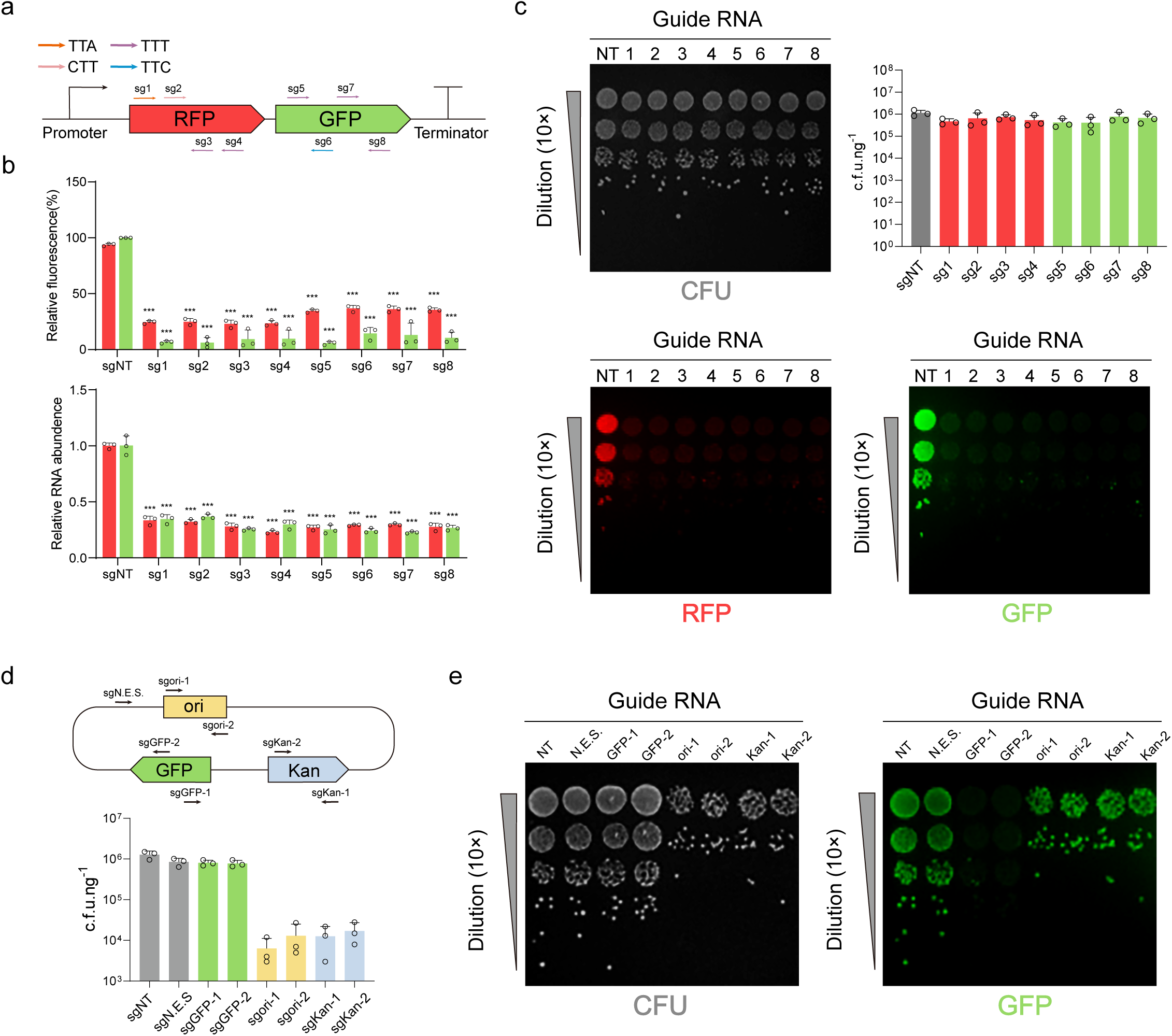
PmCas12m silences gene expression *in vivo* independent of nuclease activity. **a**, Schematic of the dual-color fluorescence interference assay using the pTarget-RFP-GFP operon reporter. The reporter encodes a bi-cistronic operon with RFP and GFP genes. Arrows denote sgRNAs designed for PmCas12m targeting. **b**, Top: Levels of RFP and GFP fluorescence in cells expressing targeting sgRNAs compared to non-targeting sgRNAs. Bottom: Relative RFP and GFP mRNA levels in cells expressing targeting sgRNAs compared to non-targeting sgRNAs. Data are presented the mean ± s.d. from three biological replicates. ***P value < 0.001. **c**, Fluorescence interference assay evaluating transformation efficiency and RFP/GFP expression *in vivo* using PmCas12m. **d-e,** Transformation assay assessing transformation efficiency with PmCas12m. Arrows denote the sgRNAs used for targeting. N.E.S., non-essential sequence.

### PmCas12m functions as an epigenome editor in human cells

Inspired by the property of PmCas12m, specifically its ability to bind target DNA without cleaving it, we propose that it could be highly suitable for epigenome editing. To test this hypothesis, we fused PmCas12m with a tripartite VP64-P65AD-Rta (VPR), which is well-established for robust transcriptional activation^34^, to explore its potential as an epigenome editing tool in human cells. The plasmid encoding the PmCas12m-VPR and the U6 promoter-driven sgRNA expression plasmid were transiently transfected into TRE3G-GFP HEK293T cells (Fig. 3a). The sgRNAs were designed to target the TRE3G promoter with TTA, TTC, or TTT PAMs, respectively (Supplementary Table 2). After 72 hours post-transfection, we measured the GFP expression via FACS. Notably, targeting the TRE3G promoter with TTA and TTC PAM sequences resulted in superior activation compared to the TTT PAM (Fig. 3a). Furthermore, human-codon-optimized PmCas12m did not exhibit higher activity than the non-codon-optimized version (Fig. 3a), therefore we selected the non-codon-optimized version for subsequent experiments. As indicated by the previous studies^35–37^, the proper positioning and arrangement of the effector domain and nuclear localization signals (NLSs) are crucial for the editing efficiency, as they impact both expression and stability. To explore this, we constructed a series of fusion variants, comparing our original PmCas12m-VPR construct (fusion#1) with eight additional variants (fusion#2-9) that feature varying arrangements of the VPR domain and NLSs (Extended Data Fig. 3). The results showed that the optimized variant (fusion #4) outperformed the others (Fig. 3b), which was chosen for further analysis.

**Fig. 3.**
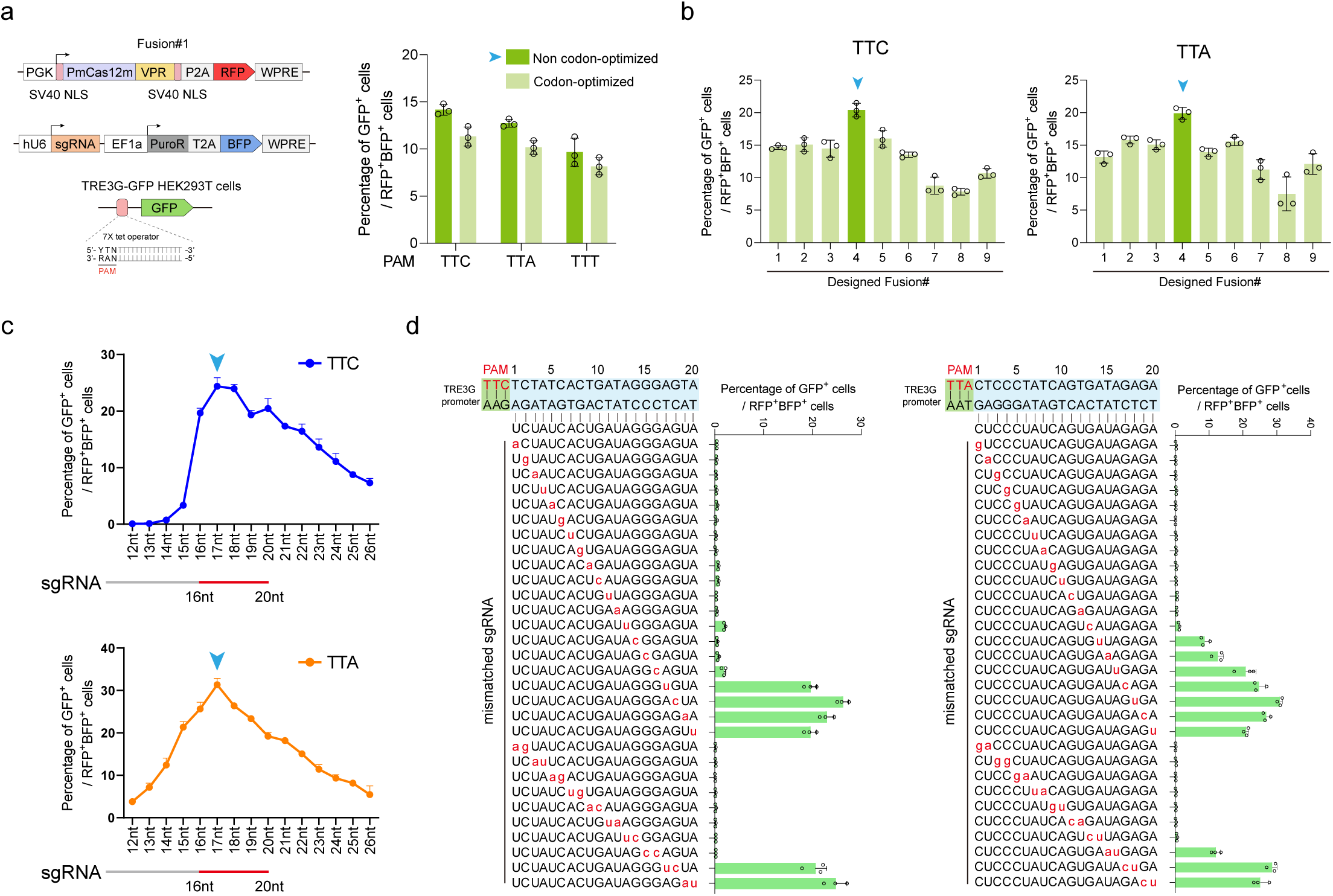
PmCas12m-mediated epigenome editing for reporter gene activation in human cells. **a**, Left: Schematic of the design strategy for the PmCas12m-VPR activation system in TRE3G-GFP HEK293T cells. Right: GFP activation efficiency of PmCas12m-VRP is measured using flow cytometry. The sgRNA targets the seven repeats in the TRE3G promoter, utilizing TTA, TTC, and TTT PAMs. **b**, GFP activation efficiency of various PmCas12m-VRP fusion designs (#1-#9) were evaluated. sgRNAs target the TRE3G promoter using TTA and TTC PAMs. **c**, PmCas12m-VPR-mediated GFP activation with different lengths of guide segments. **d**, The effects of 1-nt or 2-nt mismatches within the guide segment on PmCas12m-VPR-mediated GFP activation efficiency. Data represent the mean ± s.d. of three biological replicates.

Following the optimization of the fusion construct, we next sought to determine the effect of spacer segment length on the epigenome editing efficiency. It is well known that the length of the spacer sequence can significantly impact both the stability of the sgRNA-DNA complex and the specificity of targeting. To this end, we generated sgRNAs with varying spacer lengths ranging from 12-nt to 26-nt and evaluated their effects on transcriptional activation. Our results demonstrated that the sgRNA achieved optimal efficiency with a spacer length of 17-nt, while the efficiency declined when the spacer length exceeded 20-nt (Fig. 3c). Furthermore, to evaluate the guide – target mismatch tolerance of the PmCas12m-based epigenome editing, we designed a series of spacers containing single- or double-nucleotide mismatches with the target sequence. Our results revealed that mismatches near the PAM-proximal region—specifically, single-nucleotide mismatches within 12-nt or double-nucleotide mismatches within 14-nt of the PAM — completely abolished the epigenome editing activity, demonstrating that PmCas12m exhibits high fidelity and specificity in epigenome editing (Fig. 3d). Taken together, these results demonstrated the potential of PmCas12m as an epigenome editing tool in human cells, highlighting its applicability in precise gene regulation and epigenetic modulation.

### Cryo-EM structure of the PmCas12m-crRNA-target DNA complex

To elucidate the molecular mechanism of PmCas12m, we utilized cryo-EM to analyze the complex structure of the full-length wild-type PmCas12m with a 53-nt crRNA (including a 20-nt guide segment at the 3’ end) and 36-base pair (bp) dsDNA target with a 5’-TTA-3’ PAM sequence. The 3D reconstruction of the ternary complex was achieved at an overall resolution of 3.45 Å (Extended Data Fig. 4). The structure reveals that PmCas12m adopts a bilobed architecture, consisting of a recognition (REC) and a nuclease (NUC) lobe, connected by a linker loop (Fig. 4a-c). The REC lobe comprises the wedge (WED), REC1 and REC2 domains, while the NUC lobe is comprised of the RuvC and the target nucleic acid-binding (TNB) domains (Fig. 4a-c). The crRNA-target DNA heteroduplex is accommodated the positively charged central channel formed by the REC and NUC lobes, and recognized by PmCas12m through interactions with its sugar-phosphate backbone (Fig. 4b and c).

**Fig. 4.**
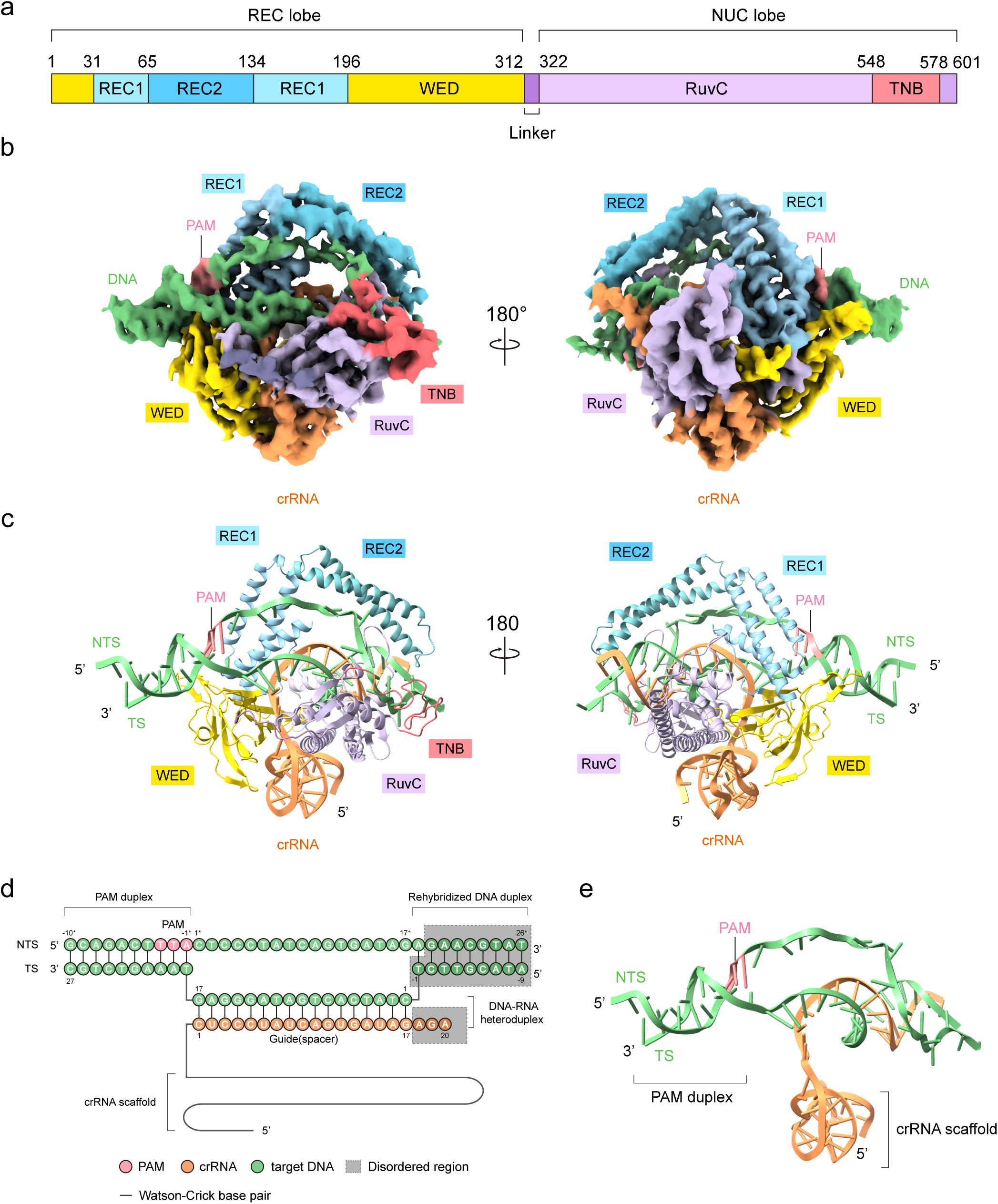
Structure of PmCas12m–crRNA–target DNA complex. **a**, Domain organization of PmCas12m. **b**, Cryo-EM map of PmCas12m–crRNA–target DNA complex. **c**, The overall structure of the PmCas12m–crRNA–target DNA complex. **d**, Schematic of the crRNA and target DNA. The disordered regions are enclosed in grey boxes. **e**, Structure of the crRNA and target DNA complex.

The crRNA structure of PmCas12m includes a 20-nt guide segment (C1 to A20) and a 33-nt crRNA scaffold (A(-33) to C(-1)) (Fig. 4d, e and Extended Data Fig. 6). In the ternary structure, nucleotides C1 to G17 in the crRNA pair with dC1 to dG17 of the target DNA strand (TS), forming a 17-bp crRNA-target DNA heteroduplex (Fig. 4d, e and Extended Data Fig. 6). This suggests that a 17-bp DNA-RNA heteroduplex represents the optimal length for PmCas12m-mediated DNA recognition, which explains our result that sgRNA achieves optimal efficiency with a spacer length of 17-nt (Fig. 3c). Within the crRNA structure, nucleotides G(-28) to C(-26) form canonical Watson-Crick base pairs with G(-15) to C(-17), while nucleotides C(-23) to C(-20) form canonical Watson-Crick base pair with G(-3) to G(-6), thereby stabilizing the crRNA scaffold (Fig. 4d,e and Extended Data Fig. 6). The crRNA is recognized by the WED and the RuvC domain, nucleotide G(-3) forms a hydrogen bond and stacking interaction with Asp475, while the backbone phosphate of G(-6) forms a hydrogen bond with Arg228 of the WED domain (Extended Data Fig. 5a, b and Extended Data Fig. 6). Additionally, nucleotides A(-29), G(-12), and C1 extensively interact with the residues of the WED and RuvC domain, including, Arg262^WED^, Arg452^RuvC^, Lys456^RuvC^, Arg457^RuvC^ and Lys540^RuvC^ (Extended Data Fig. 5a, c, e and Extended Data Fig. 6). In addition, backbone phosphate groups of A12 and G13 in TS interact with Try528 and Arg499, respectively (Extended Data Fig. 5a, d and Extended Data Fig. 6). In the structure of the PAM duplex, NTS sequences of the 5′-TTA-3′ PAM interact with the REC1 domain. The phosphate backbone of dT(-3*) and dT(-2*) forms hydrogen bonds with Try166 and Arg156, respectively, while the nucleotide of dT(-2*) also interacts with Asn167 in the REC1 domain (Extended Data Fig. 5a, f and Extended Data Fig. 6). These structural features explain the recognition mechanism of the 5’-TTA-3’ PAM sequence for PmCas12m-mediated DNA binding.

Unlike most Cas12 enzymes, where the RuvC active site contains conserved DED catalytic residues for target dsDNA cleavage, PmCas12m exhibits a non-canonical motif. The conserved DED residues are replaced by Asn372, Asp494, and Asp581 within the RuvC domain, respectively (Extended Data Fig. 5a, g). Furthermore, nucleotides dG(11*) to dG(13*) in the NTS are located at this non-canonical NDD RuvC active site (Extended Data Fig. 5a, g). A previous study on MmCas12m demonstrated that a non-canonical HDD motif leads to the absence of the second magnesium ion, resulting in the loss of cleavage activity for the target DNA^33^. Therefore, we propose that the NDD motif in PmCas12m exhibits similar characteristics, potentially explaining its lack of nuclease activity. Overall, the cryo-EM structure of the PmCas12m-crRNA-target DNA complex provides critical insights into the molecular mechanism of PmCas12m, particularly its substrate recognition and interaction.

### Structure-guided engineering of PmCas12m for enhanced gene activation in human cells

To augment the epigenome editing activity of this miniature protein, we employed DMS to investigate how amino acid substitutions affect the editing efficiency in HEK293T cells. Through analysis of the PmCas12m-crRNA-target DNA structure, we selected 137 residues based on their accessibility and proximity to the binding interfaces with both the crRNA and target DNA (Fig. 5a, c). These residues were divided into four units: residues 121-185, residues 226-264, residues 306-315, and residues 493-515 (Fig. 5c). For each unit, separate PmCas12m-VPR libraries were constructed, each encompassing all 20 possible amino acid substitutions (Fig. 5c). Subsequently, sgRNAs targeting TRE3G promoter with TTA or TTC PAMs were transduced into TRE3G-GFP HEK293T cells. The PmCas12m-VPR variant libraries were packaged into lentivirus and transduced at a multiplicity of infection (MOI) of less than 0.3, ensuring that no more than one PmCas12m-VPR mutant was expressed per cell. 72 hours post-infection, cells with high GFP expression and low GFP expression were sorted via FACS and cultured for an additional 5 days. After 5 days, we re-sorted the cells to prevent contamination from cells of other populations. Total RNA was then extracted from high GFP-expression cells and low GFP-expression cells, and deep sequencing analysis was performed, using the low GFP-expression cells served as the control (Fig. 5b). DMS experiments were conducted using two different sgRNAs targeting distinct PAM sequences, which yielded similar results (Fig. 5c, d and Extended Data Fig. 7).

**Fig. 5.**
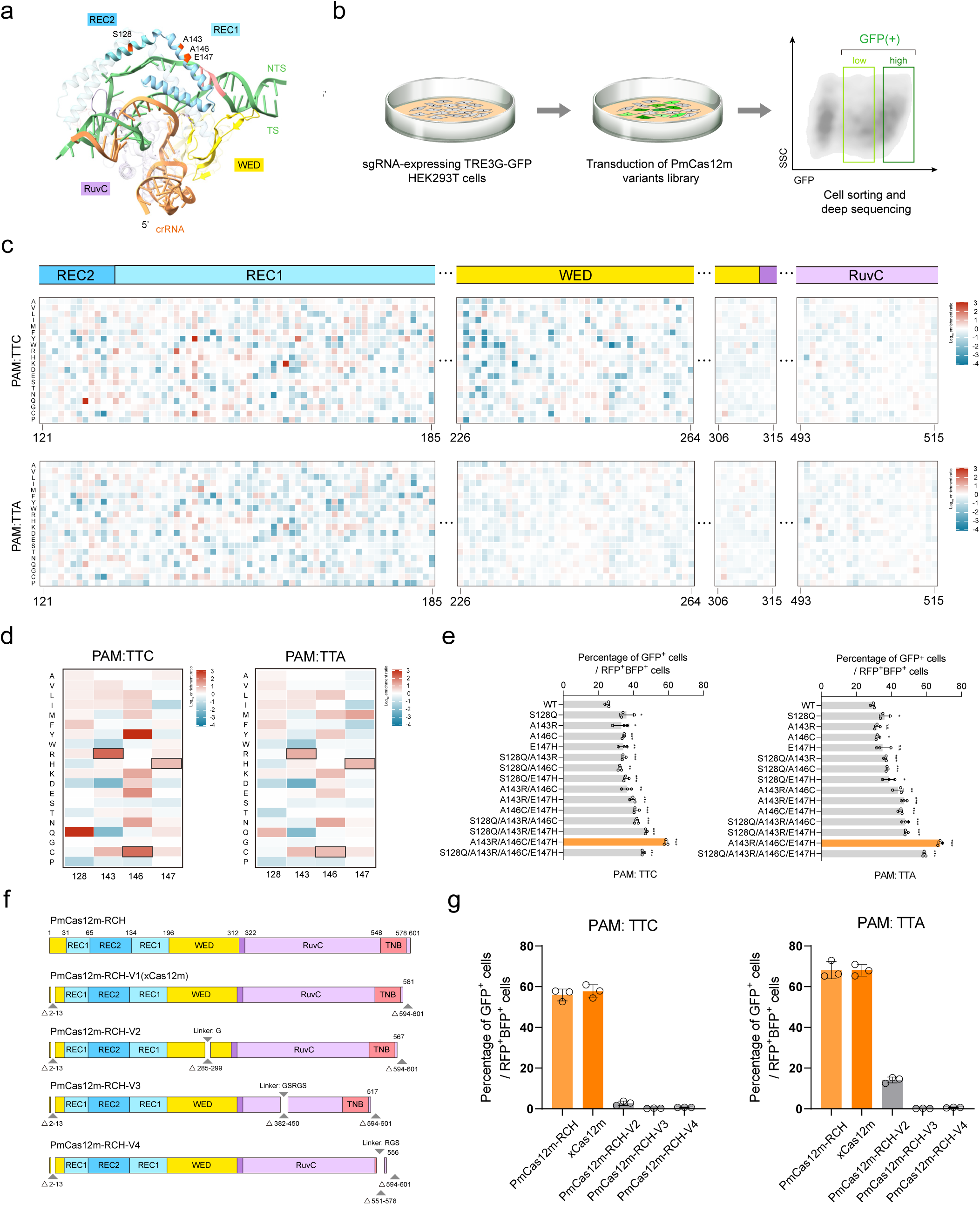
Iterative protein engineering of PmCas12m variants with deep mutational scanning. **a**, The overall structure of the PmCas12m-crRNA-target DNA complex serves as the basis for structure-guided engineering of PmCas12m. Amino acid residues selected for structure-guided rational design are highlighted in their respective colors. **b**, Schematic of the DMS method to evaluate the GFP activation efficiency in TRE3G-GFP HEK293T cells. **c**, Heatmap illustrating the effects of all single mutations on GFP activation efficiency with TTC (top) and TTA (bottom) PAMs. Colors represent the mutational effects based on the scale bars, with red indicating more efficient mutations. **d**, Close-up view of the key amino acid substitutions with TTC (left) and TTA (right) PAMs. Mutations selected for PmCas12m-RCH are highlighted with gray-colored borders. **e**, GFP activation efficiency of PmCas12m-VPR variants was measured by flow cytometry with TTC (left) and TTA (right) PAMs. Data represent the mean ± s.d. of three biological replicates. ***P value < 0.001, **P value < 0.01, *P value < 0.05, n.s., non-significant. **f**, Schematic of PmCas12m with rational deletions according to the structure analysis. **g**, GFP activation activity of PmCas12m deletion variants with TTC (left) and TTA (right) PAMs. Data represent the mean ± s.d. of three biological replicates.

Based on the structural analysis and DMS results, four mutations (S128Q, A143R, A146C, and E147H) stood out, which were located near the nucleic acids and showed a higher epigenome editing efficiency compared to the control (Fig. 5c, d and Extended Data Fig. 7). Using TRE3G-GFP HEK293T cells, we observed a slight enhancement in activation for all four variants (Fig. 5e). To further improve the activity of this compact protein, we performed a second round of optimization. In this round, each of the four mutated residues was individually combined to create six double-mutant variants, all of which demonstrated higher activity compared to the first round (Fig. 5e). The third round of screening included three triple variants and one quadruple variant based on the second-round results. Among these, the A143R/A146C/E147H variant exhibited the highest performance compared to wild-type PmCas12m, which was named PmCas12m-RCH (Fig. 5e). Together, we showed that the PmCas12m mutants with A143R/A146C/E147H substitutions exhibited enhanced epigenome editing activity.

Epigenome editing tools require the fusion of various effector domains, resulting in a relatively large overall size, even when using a compact protein like PmCas12m-RCH. This large size presents challenges for delivery efficiency. Therefore, rational deletions of PmCas12m-RCH could significantly enhance the delivery efficacy, facilitating the development of effective epigenome editing therapies. To empirically determine the optimal size range for functional deletions, we analyzed the structure of PmCas12m and identified five potential deletion regions, which are either flexible or not involved in protein-DNA/RNA binding. Intriguingly, PmCas12m-RCH with both N-terminal (Δ2-13) and C-terminal (Δ594-601) deletions retained near PmCas12m-RCH levels of epigenome editing efficiency, which we name as PmCas12m-RCH-V1 (Fig. 5f, g). However, when we introduced additional deletions individually in the WED (Δ285-299), RuvC (Δ382-450), or TNB (Δ551-578) regions of PmCas12m-RCH-V1 to further evaluate its function, we found that these deletion variants exhibited a marked reduction or complete loss of editing activity (Fig. 5f, g). Taken together, through iterative protein engineering and screening, we established an efficient PmCas12m variant with deletions in the N-terminal (Δ2-13) and C-terminal (Δ594-601) regions of PmCas12m-RCH, resulting in a compact 581-amino-acid protein (Fig. 5f, g), which we refer to simply as “xCas12m” hereafter.

### xCas12m enables highly potent and specific epigenome editing

To assess the application potential of xCas12m, we evaluated its epigenome editing activity and compared it with dCas9^4^, dSpRY^38^, wild-type PmCas12m and four compact nuclease-dead CRISPR-Cas systems (referred to as dCasMINI^37^, MmCas12m^13^, denAsCas12f1^39^ and dSpCas12f1^40^). To minimize the chromatin context interference arising from targeting different genomic loci, sgRNAs for each system were carefully designed to target the sequences within a narrowly overlapping range (Supplementary Table 2). To evaluate the epigenome activation efficiency of xCas12m-VPR (Fig. 6a), we measured the expression levels of seven endogenous genes in HEK293T cells using qRT-PCR. For each gene, sgRNAs were specifically designed to target promoter regions proximal to the transcriptional start site (TSS), enabling precise modulation of gene activation (Supplementary Table 2). The results indicated that both wild-type PmCas12m-VPR and xCas12m-VPR efficiently and robustly induced activation across these endogenous genes, with xCas12m-VPR consistently outperforming wild-type PmCas12m-VPR in activation efficiency (Fig. 6b-h), showcasing the effectiveness and practicality of our protein engineering approach in enhancing the epigenome editing activation. Importantly, xCas12m-VPR demonstrated superior activation potential compared to the widely used dCas9-VPR in certain genes (Fig. 6b-d), highlighting its promising capabilities for epigenome editing applications.

**Fig. 6.**
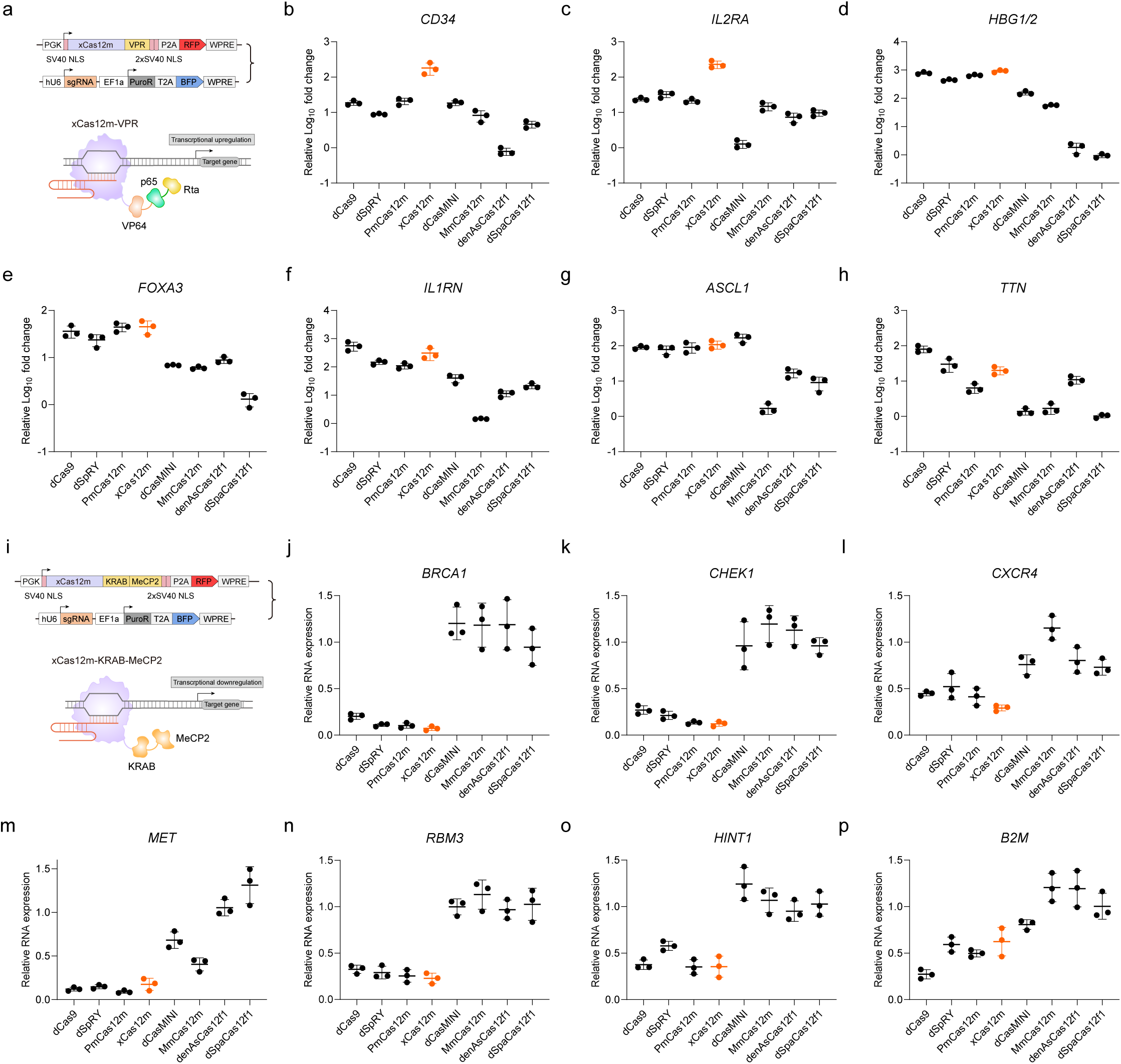
Endogenous gene regulation using xCas12m-medicated epigenome editing in human cells. **a**, Schematic of constructs used for the xCas12m-VPR activation system. **b-h**, Comparison of gene activation efficiency using xCas12m-VPR, PmCas12m-VPR, dCas9-VPR, dSpRY-VPR, MmCas12m-VPR and three compact dCas12f-VPR systems when targeting the promoters of *CD34*(**b**), *IL2RA*(**c**), *HBG1/2*(**d**), *FOXA3*(**e**), *IL1RN*(**f**), *ASCL1*(**g**) and *TTN*(**h**). Data represent the mean ± s.d. of three biological replicates. **i**, Schematic of constructs used for the xCas12m-KRAB-MeCP2 repression system. **j-p**, Comparison of gen repression efficiency using xCas12m-KRAB-MeCP2, PmCas12m-KRAB-MeCP2, dCas9-KRAB-MeCP2, dSpRY-KRAB-MeCP2, MmCas12m-KRAB-MeCP2 and three compact dCas12f-KRAB-MeCP2 systems when targeting the promoters of *BRCA1*(**j**), *CHEK1*(**k**), *CXCR4*(**l**), *MET*(**m**), *RBM3*(**n**), *HINT1*(**o**) and *B2M*(**p**). Data represent the mean ± s.d. of three biological replicates.

Given the immense potential of xCas12m for CRISPRa, we sought to further investigate its broader applicability in epigenome editing by assessing whether it could also downregulate endogenous target genes. To achieve this, we employed the well-established repressor KRAB-MeCP2, an effector commonly used for CRISPR-based gene silencing^41^, and fused it with the eight CRISPR-Cas systems mentioned above (Fig. 6i). As with the activation experiments, sgRNAs were designed to target promoter sequences near the TSS to ensure consistency and precision in gene modulation (Supplementary Table 2). The results of xCas12m-KRAB-MeCP2 revealed varying degrees of gene repression across all seven endogenous genes. Importantly, xCas12m-KRAB-MeCP2 exhibited efficient repression activity, demonstrating gene silencing effects that were comparable to those observed with dCas9-KRAB-MeCP2 and its variants dSpRY-KRAB-MeCP2 (Fig. 6j-p). In several cases, xCas12m-KRAB-MeCP2 even showed higher repression efficiency than dCas9-KRAB-MeCP2 (Fig. 6j-l), suggesting that xCas12m-KRAB-MeCP2 could be an effective alternative for CRISPRi approach.

CRISPR-based systems can induce off-target effects or cause unintended epigenetic changes^2,3^. To evaluate the specificity of xCas12m-VPR-mediated epigenome editing, we targeted the *HBG1/2* promoter and performed whole-transcriptome RNA sequencing (RNA-seq) to assess the potential off-target effects on a genome-wide scale. The results revealed that xCas12m-VPR showed the strongest activation signal for *HBG1* and *HBG2* (Extended Data Fig. 8a). Furthermore, ChIP-seq analysis revealed significant enrichment of xCas12m-VPR at the targeted *HBG1* and *HBG2* promoters, with minimal off-target binding compared to the non-targeting control (Extended Data Fig. 8b). Due to the high sequence homology between the *HBG1* and *HBG2* promoters, the sgRNA designed for one could not distinguish between the two loci, complicating the assignment of differential effects solely to one gene. Except for the few differentially expressed genes, we did not observe any significant off-target transcriptional changes in neighboring genes, and none of the differentially expressed genes exhibited a near-sequence match to the *HBG1/2* promoter-targeting sgRNA. This suggested that the observed gene expression changes were not the result of improper targeting by the epigenome editor but rather reflect the intended, locus-specific modulation of the gene activity. Taken together, these results underscore the potential of xCas12m as an effective and highly adaptable platform for precise epigenome editing, paving the way for future studies and applications in genetic research and therapeutic gene regulation.

### xCas12m-CRISPRoff efficiently suppresses HBV transcription and replication

To investigate the efficiency of xCas12m-mediated inhibition of HBV infection, we developed xCas12m-CRISPRoff (Fig. 7a), an epigenome editing tool based on the CRISPRoff strategy^42^, aiming at offering a potentially more efficient and scalable approach to silence HBV transcription and replication. We evaluated its performance using three distinct HBV cell models: (1) HepG2 cells transfected with 1.3× HBV plasmids, (2) the HepAD38 cell line, which integrates the HBV genome and expresses HBV under tetracycline control, and (3) HBV-infected HepG2-NTCP cells. Nine sgRNAs were designed to target four viral promoters and two viral enhancers, aiming at key regulatory elements critical for HBV transcription and replication (Fig. 7b and Supplementary Table 2). Firstly, we evaluated the antiviral efficacy by measuring HBeAg and HBsAg levels in the culture supernatant at the indicated time points. As expected, all sgRNA candidates resulted in the reduction of HBeAg and HBsAg levels in these HBV cell models, although the efficiency of interference varied across different sgRNAs (Fig. 7c, Extended Data Fig. 9a and b). Secondly, with potent sgRNAs, we assessed the antiviral efficacy by quantifying the levels of HBV total RNA and pgRNA, since HBV total RNA serves as an indicator of overall viral transcription, whereas pgRNA functions as a crucial intermediate in the replication of the viral genome. A consistent reduction in both total RNA and pgRNA levels was observed, indicating that xCas12m-CRISPRoff effectively interfered with HBV transcription and replication (Fig. 7d, Extended Data Fig. 9c and d). Similarly, RNA-seq analysis demonstrated that xCas12m-CRISPRoff significantly reduced the abundance of HBV transcripts in HBV-infected HepG2-NTCP cells (Fig. 7e). Furthermore, xCas12m-CRISPRoff precisely targeted the HBV genome, with minimal impact on the human transcriptome (Extended Data Fig. 10). Thirdly, we examined both total HBV DNA and covalently closed circular DNA (cccDNA) levels in HBV-infected HepG2-NTCP cells, considering total HBV DNA represents the overall viral genome, while cccDNA serves as the episomal form of the HBV genome, both of which are critical for viral DNA replication. As expected, xCas12m-CRISPRoff significantly reduced the levels of both total HBV DNA and cccDNA in the HBV-infected HepG2-NTCP cell model (Fig. 7f, g), suggesting that xCas12m-CRISPRoff effectively interfered with HBV genome maintenance.

**Fig. 7.**
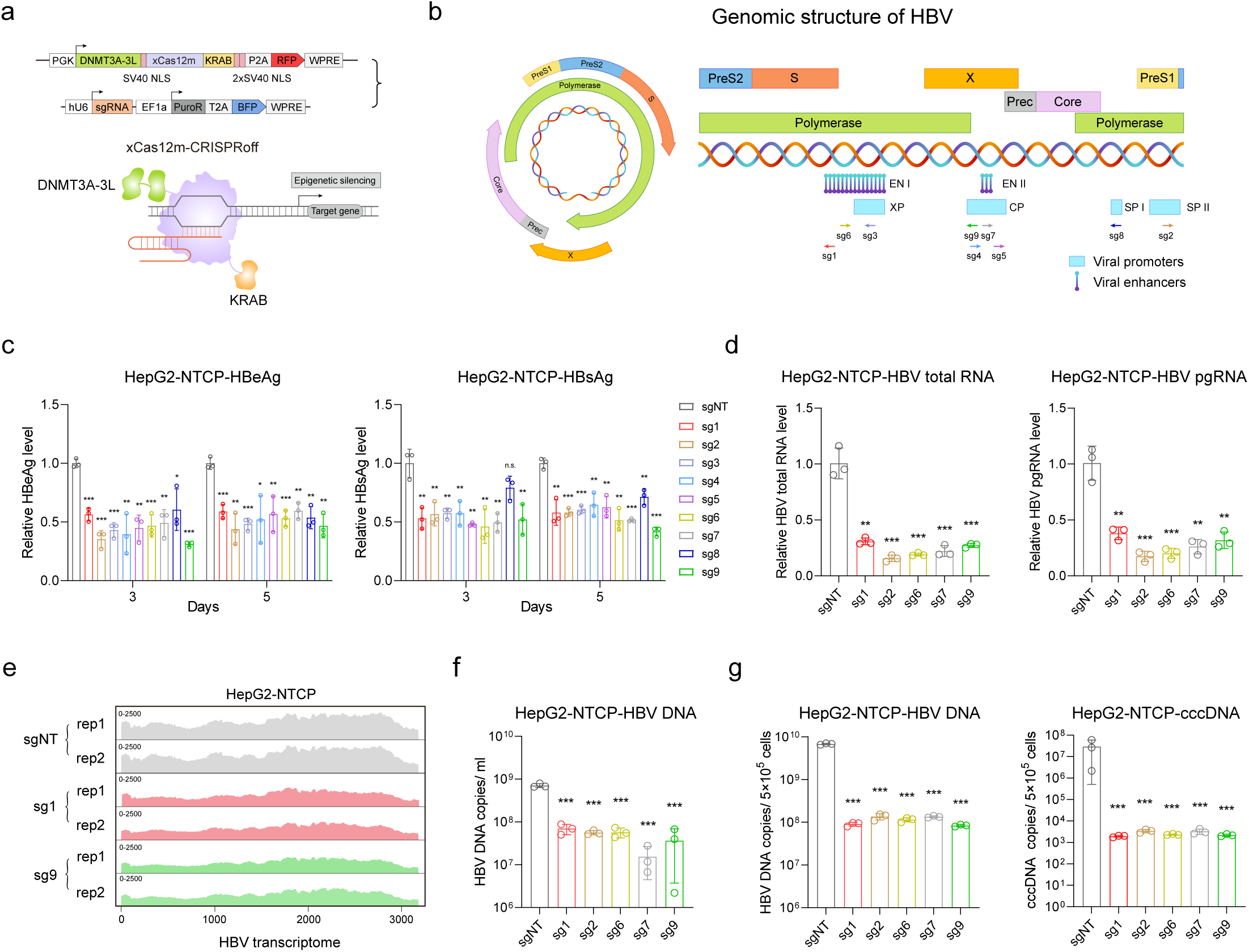
xCas12m-CRISPRoff-medicated epigenetic editing in HBV-infected cell models. **a**, Schematic of constructs used for the xCas12m-CRISPRoff repression system. **b**, Circular(left) and linear(right) diagrams of the HBV genome. XP, X promoter; CP, Core promoter; SP I, PreS1 promoter; SP II, PreS2 promoter; EN I, Enhancer I; EN II, Enhancer II. sgRNAs tested for targeting four viral promoters and two viral enhancers within the HBV genome. **c**, Levels of HBeAg (left) and HBsAg (right) in cell culture supernatant of HBV-infected HepG2-NTCP cells at the indicated time points post-transfection with xCas12m-CRISPRoff systems. Data represent the mean ± s.d. of three biological replicates. ***P value < 0.001, **P value < 0.01, *P value < 0.05, n.s., non-significant. **d**, Levels of HBV total RNA (left) and pgRNA (right) in HBV-infected HepG2-NTCP cells on the 5th day post-transfection with xCas12m-CRISPRoff systems. Data represent the mean ± s.d. of three biological replicates. ***P value < 0.001, **P value < 0.01. **e**, RNA-seq profiles showing the HBV transcriptome in HBV-infected HepG2-NTCP cells upon xCas12m-CRISPRoff-mediated repression of critical viral elements using two representative sgRNAs. Independent replicate experiments are shown as rep1 and rep2, respectively. **f**, Total HBV DNA levels in cell culture supernatant of HBV-infected HepG2-NTCP cells on day 5 post-transfection with xCas12m-CRISPRoff systems. Data represent the mean ± s.d. of three biological replicates. ***P value < 0.001. **g**, Levels of total HBV DNA (left) and cccDNA (right) in HBV-infected HepG2-NTCP cells on day 5 post-transfection with xCas12m-CRISPRoff systems. Data represent the mean ± s.d. of three biological replicates. ***P value < 0.001.

Having determined the antiviral efficacy of xCas12m-CRISPRoff *in vitro*, we further evaluated its therapeutic potential in the AAV-HBV1.04 mouse model^43^. Given the compact size of xCas12m-CRISPRoff (1305 amino acids), we successfully engineered the xCas12m-CRISPRoff system, along with its sgRNA components, into a single AAV vector (Fig. 8a). This configuration is well-suited to the packaging limitations of AAV vector, potentially facilitating effective *in vivo* delivery for potential treatment of HBV infection. To assess whether xCas12m-CRISPRoff exhibited potent anti-HBV activity *in vivo*, we established an HBV mouse model by injecting recombinant AAV-HBV1.04 using AAV serotype 8 (AAV8). After 1 week, mice were randomly assigned to three groups and received tail vein injection with AAV-xCas12m-CRISPRoff-sgNT, AAV-xCas12m-CRISPRoff-sg1 or AAV-xCas12m-CRISPRoff-sg9, respectively. Blood samples were subsequently collected at the indicated time points (Fig. 8a). The results showed that xCas12m-CRISPRoff significantly reduced the serum HBeAg levels, HBsAg levels and circulating HBV DNA levels (Fig. 8b-d). Notably, we observed that the reduction in viral products was durable following a single dose (Fig. 8b-d), attributable to the heritable gene silencing memory of CRISPRoff. Overall, these findings underscore the potential of xCas12m-CRISPRoff as a promising epigenome editing tool for inhibiting HBV antigen expression and circulating HBV DNA, thus providing a new avenue for the development of epi-silencing therapies against HBV infection.

**Fig. 8.**
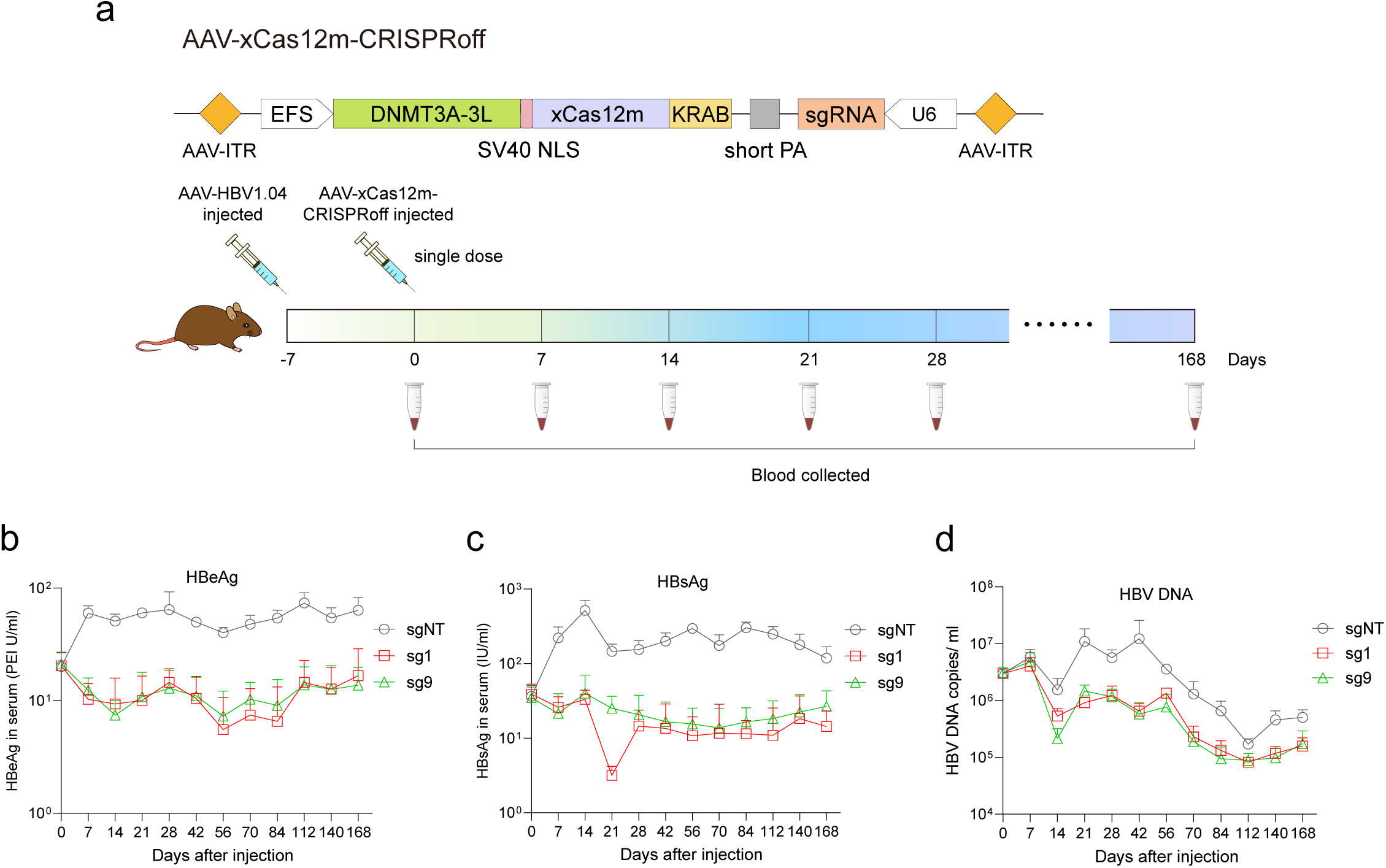
xCas12m-CRISPRoff-medicated epigenetic silencing for HBV treatment. **a**, Schematic of AAV-based all-in-one xCas12m-CRISPRoff repression system (top) and *in vivo* experimental design to evaluate the efficiency of AAV-based xCas12m-CRISPRoff system for HBV treatment (bottom). **b-d**, Levels of HBeAg (**b**), HBsAg (**c**) and circulating HBV DNA (**d**) in mouse serum at the indicated time points. Data represent the mean ± s.d. of 5 mice in each group.

## Discussion

In this study, we demonstrate that subtype V-M CRISPR-Cas protein, PmCas12m, is a programmable RNA-guided DNA-binding protein that lacks the conserved DED residues in the RuvC domain. This observation is particularly intriguing, as previous analyses of catalytically active Cas12 and their ancestral TnpB proteins have shown that the RuvC domain typically contains a highly conserved DED motif, which is crucial for endonuclease activity^44–46^. However, within the subtype V-M CRISPR-Cas system, these residues exhibit considerable variability, highlighting the structural and functional diversity across other subtypes of type V CRISPR-Cas systems^12,13^. TnpB proteins have independently evolved on multiple occasions throughout evolutionary history, leading to the emergence of diverse CRISPR-Cas12 subtypes that retain the RuvC DED active site^47^. By contrast, the absence of these conserved residues in PmCas12m raises intriguing questions about the evolutionary trajectory of subtype V-M CRISPR-Cas proteins. Interestingly, a recent study has identified a novel class of TnpB-derived proteins, called TldRs, which also lack conserved DED residues in the RuvC domain and function as naturally occurring transcriptional factors^14^. Based on these findings, we propose that TldRs may represent the ancestral form of the subtype V-M CRISPR-Cas system, providing a potential explanation for the functional divergence observed from canonical type V CRISPR-Cas proteins.

Given that PmCas12m robustly binds to dsDNA without nuclease activity, it is worth considering whether this protein can still provide defense against invading MGEs. CRISPR-Cas systems have evolved across diverse microbial species to provide adaptive immunity against invading phages and other MGEs, such as plasmids and transposons^48^. Immunity conferred by class 2 type V CRISPR-Cas systems typically relies on the RuvC nuclease domain to recognize and cleave foreign genetic elements^46,49^. However, recent studies have revealed that Cas12 homologs, specifically Cas12c and Cas12m, can interfere with MGEs through targeted DNA binding, indicating alternative modes of interference that impede either the replication of MGE genomes or the transcription of essential MGE genes^9,12,13^. These findings demonstrate that the immune capabilities of type V CRISPR-Cas systems extend beyond simple nuclease-mediated destruction, incorporating diverse mechanisms to counteract invading genetic elements. Building on this understanding, it is unsurprising that PmCas12m also employs an interference mechanism through RNA-guided binding to complementary dsDNA targets in MGEs. This characteristic explains the biological significance of PmCas12m in nature as an alternative defense strategy against invading genetic elements. Furthermore, its mechanism of action opens new possibilities for leveraging this protein in various applications, such as epigenome modulation, transcriptional regulation, base editing and other precision biotechnological applications.

Currently, the most widely used tools for epigenome editing are based on dCas9 and its variants. By fusing these systems with diverse effector domains, researchers can precisely modulate the epigenetic landscape at specific genomic loci, enabling the regulation of gene expression and cell fate determination. Despite their potential, translating these capacities into clinical applications remains a significant challenge. AAV-based vectors have become increasingly popular for therapeutic purposes due to their safety and ability to transduce a wide range of cell types. Nevertheless, the large size of dCas9-based systems exceeds the packaging capacity of AAV vectors (∼4.7 kb), restricting their effective delivery^6,7^. To address this limitation, we engineered a miniaturized version of the PmCas12m, termed xCas12m, which consists of 581 amino acids and demonstrates high specificity and efficiency in epigenome editing. At approximately 40% the size of spCas9, xCas12m’s hypercompact size is particularly well-suited for fusion with highly functional but larger effectors in AAV-based gene delivery applications. Additionally, its miniature size provides considerable potential for further optimization, including reducing its size even further, enhancing its activity, and engineering the domains to improve editing efficiency and specificity. Such optimizations could further expand the potential of xCas12m as a robust platform for epigenome editing. Beyond its smaller size, xCas12m offers a distinct advantage due to its PAM-flexible recognition ability. This feature allows it to target regions of the genome that may be less accessible to conventional CRISPR/Cas systems, broadening its potential applications in both research and therapeutic contexts. Taken together, our data suggest that xCas12m-based epigenome editors could be particularly advantageous in contexts where larger CRISPR/dCas systems face delivery challenges. Its compact size, efficiency, and deliverability make it a highly promising candidate for versatile and efficient epigenome editing, with significant potential for applications requiring precise, scalable, and effective gene regulation.

The HBV genome is a 3.2 kb relaxed-circular DNA (rcDNA) with a complete minus strand and an incomplete plus strand. Upon infection, the rcDNA is transported to the host cell nucleus, where it is converted into cccDNA through host DNA repair mechanisms. In addition to cccDNA formation, a small fraction of rcDNA is converted into longer double-stranded linear DNA (dslDNA), which can integrate into the host genome^50–53^. Given these complexities, altering the epigenetic regulation of infected cells represents a crucial strategy for developing therapies aimed at eliminating or at least achieving complete inactivation of the HBV genome^26,27^. A recent study has demonstrated that the CRISPRoff system can achieve stable and long-term repression of gene expression by modifying the epigenetic landscape, providing a promising strategy for sustained gene silencing^42^. Building on this concept, we developed a novel epigenome editing tool, xCas12m-CRISPRoff, which incorporated the compact xCas12m with the CRISPRoff strategy. The compact size of xCas12m-CRISPRoff (1305 amino acids) enables it to be packaged alongside sgRNA into a single AAV vector, effectively overcoming the packaging constraints and significantly enhancing delivery efficiency. This streamlined design enhances the practicality of xCas12m-CRISPRoff for *in vivo* applications, offering a robust solution for efficient and precise epigenetic modulation. The efficacy of this approach was validated in an HBV-infected mouse model, where the xCas12m-CRISPRoff system exhibited a robust ability to suppress HBV antigens and circulating HBV DNA with a single dose, demonstrating the stability and durability of epigenetic silencing. These results highlight the potential of xCas12m-CRISPRoff to achieve long-term viral suppression through precise epigenetic silencing, providing a distinct advantage over traditional antiviral therapies. Based on these findings, the xCas12m-CRISPRoff platform represents a significant advancement in epigenetic therapy, with the potential to revolutionize HBV treatment and extend its applicability to other diseases requiring precise and durable epigenetic remodeling. Nonetheless, considering the presence of rcDNA, cccDNA, and integrated HBV DNA in HBV-infected cells, further investigation is warranted to evaluate the specific efficacy of xCas12m-CRISPRoff in addressing these distinct forms of the HBV genome. Beyond HBV, xCas12m-mediated epigenetic editing platform holds broad potential for application in other infectious diseases caused by DNA-based pathogens. By leveraging its advantage of compact size, high specificity, and the heritable and reversible features of native epigenetic regulation to modulate gene expression without introducing DNA breaks, xCas12m-based epigenome editing tools can be adapted for therapeutic applications across a broad spectrum of viral and bacterial pathogens. For instance, persistent viruses such as human papillomavirus (HPV), Epstein-Barr virus (EBV), or herpes simplex virus (HSV), as well as bacterial pathogens that rely on epigenetic mechanisms for virulence or antibiotic resistance, may also be amenable to therapeutic targeting using this system. Moreover, the modular and programmable properties of xCas12m-based tools further support their potential for multiplexed and pathogen-specific epigenetic modifications, offering a powerful strategy for the development of anti-infective therapies.

Highly efficient and accurate protein structure prediction is critical for protein discovery and engineering. The recently developed AI methodologies, like AlphaFold 3^30^ and RoseTTAFold^54^, have achieved quick and accurate structure prediction based on limited information, potentially facilitating large-scale mining and classification of proteins with specific functions. In this study, we developed and applied a structure-guided search pipeline to identify, characterize and engineer a novel miniature xCas12m, emphasizing the power of structure-guided protein mining. In principle, this strategy has the potential to be applied to the high-throughput classification and functional analysis of any protein dataset. We envision that artificial intelligence has the potential to expedite the development process by accurately identifying optimal editing targets, enhancing CRISPR’s precision and efficiency, and minimizing off-target effects in the future. Indeed, a recent study has demonstrated the first successful precision editing of the human genome with a programmable gene editor designed entirely with machine learning, accentuating the importance of language models in the future design of Cas proteins^55^.

In conclusion, we developed and applied a structure-guided search pipeline to identify, characterize and engineer miniature xCas12m protein for highly potent and specific epigenome editing *in vitro* and *in vivo*. These findings establish xCas12m as a compact, durable, and clinically actionable tool for epigenome editing, offering a paradigm to combat diverse diseases including viral infections through precision epigenetic regulation.

## Contributions

Conceptualization: KL; Methodology: TY, MJ, DY, ZG, RZ, YJ, ZY, LQ, JM, FM, KX, LB; Writing and editing: TY, LB and KL; Supervision: KL.

## Acknowledgments

We deeply thank Dr. Chase L. Beisel and Tatjana Achmedov (Helmholtz Institute for RNA-based Infection Research, Germany) for providing the pPAM-SCANR plasmid and E. coli BW25113 competent cells. This study was supported by grants from the National Natural Science Foundation of China (KL, No. 82273131; LB, No. 32171212 and KX, No. 81873579), the National Key Research and Development Program of China (KL, No. 2021YFC2302300), the Fundamental Research Funds for the Central Universities (KL, No. BMU2021YJ073), the Clinical Medicine Plus X - Young Scholars Project of Peking University, the Fundamental Research Funds for the Central Universities (KL, No. PKU2024LCXQ044), Peking University Health Science Center Fund (KX, No. BMU2023YJ016), and Young Elite Scientists Sponsorship Program by BAST (KX, No. QNRCNRC001).

## Declarations

The authors declare no competing interests.

## Availability of supporting data

The data supporting this study are available on request from the corresponding author.

## Materials and methods

### Cell culture, transfection, and flow cytometry analysis

HEK293T, TRE3G-GFP HEK293T, HepG2, HepAD38 and HepG2-NTCP cells were cultured at 37°C with 5% CO_2_ in Dulbecco’s Modified Eagle Medium (DMEM, Gibco) supplemented with 10% fetal bovine serum (FBS, TransGen) and 1% penicillin-streptomycin (TransGen). All cell lines were tested for *Mycoplasma* contamination. For TRE3G-GFP HEK293T cells, transfections were performed with PEI (Polysciences) at a ratio of 2 μl of reagent per microgram of plasmid in 24-well plates unless otherwise noted. For each well on the plate, 600ng of corresponding Cas12m plasmids and 400ng of sgRNA plasmids were mixed with 50 μl OptiMEM Reduced Serum Medium (Gibco). Separately, 50 μl OptiMEM was mixed with 2 μl PEI reagent. The plasmid and PEI mixtures were then combined, incubated at room temperature for 20 min, and then added to the cells. The transfected cells were harvested 3 days post-transfection for flow cytometry analysis to evaluate the GFP activation efficiency. For HepG2, HepAD38 and HepG2-NTCP cells, transfections were performed with Lipofectamine 3000 (Invitrogen) according to the manufacturer’s protocol. Transfected cells were collected at the indicated times for further analysis.

### PAM-SCANR assay

The PAM-SCANR assay was performed as previously reported with the following modifications^32^. Briefly, 100ng of the PAM library plasmid and 200ng CRISPR/Cas12m plasmid were electroporated into *E. coli* BW25113 strain lacking the lacI, lacZ genes as well as the type I-E CRISPR-Cas system. After recovery, the transformation mixture was inoculated into 5 mL of LB medium (1:100 dilution) supplemented with 50 μg/mL kanamycin and 34 μg/mL chloramphenicol, and incubated overnight at 37 °C. The next day, the cultures were diluted 1:100 in PBS, and GFP-positive and GFP-negative cells were sorted using FACS. Approximately 100,000 single cells were sorted per experiment, cultured in 5mL of LB medium with the appropriate antibiotics, and incubated overnight at 37 °C. The following day, the GFP-positive sample and GFP-negative sample were diluted 1:100 in PBS and sorted separately for GFP-positive and GFP-negative cells with 100,000 single cells. The sorted cells were cultured in 5 mL of LB medium with appropriate antibiotics and incubated overnight at 37 °C. Plasmids were extracted using EndoFree Mini Plasmid Kit (TIANGEN). Approximately 200 bp amplicons surrounding the target loci were amplified with FastPfu Fly PCR SuperMix (TransGen) to add the Illumina adaptors and barcodes for each sample. The libraries were subjected to Illumina HiSeq sequencing (PE150). The abundance of each PAM was counted and normalized to the total number of reads in the corresponding sample. PAMs with an enrichment greater than 16-fold were selected to generate PAM logos^56^ and PAM wheels^57^.

### Small RNA-seq and analysis

Total RNA was extracted from *E. coli* cells harboring the CRISPR/Cas12m system using the *TransZol* Up RNA Extraction Kit (TransGen). Approximately 60ug of the extracted RNA was treated with 2 U of DNase I (NEB) at 37 °C for 30 minutes and subsequently purified by phenol/chloroform extraction. The DNA-free RNA was then incubated with 20 U of T4 polynucleotide kinase (T4 PNK, NEB) at 37 °C for 6 hours. Subsequently, 1 mM adenosine triphosphate (ATP; NEB) was added, and the reaction was incubated at 37 °C for an additional hour. After phenol/chloroform extraction, the RNA was treated with 5 U of RppH (NEB) at 37 °C for 1 hour, followed by a final phenol/chloroform purification. 5 μg of the pre-treated RNA was used to construct a small RNA library using the VAHTS Small RNA Library Prep Kit for Illumina (Vazyme), following the manufacturer’s protocol. The small RNA library was subjected to Illumina HiSeq sequencing (PE150). The sequencing data were analyzed by Bowtie2^58^ and SAMtools^59^.

### PAM depletion assay

PAM depletion assay was performed as previously described^60^. Briefly, 100ng of the PAM library and 200ng CRISPR/Cas12m plasmid were electroporated into *E.coil* DH5α. Approximately 100,000 colonies were collected to ensure sufficient coverage of all possible PAMs. Plasmids were then extracted using EndoFree Mini Plasmid Kit (TIANGEN). To prepare for sequencing, approximately 200 bp amplicons surrounding the target loci were amplified with FastPfu Fly PCR SuperMix (TransGen) to add Illumina adaptors and barcodes. The libraries were subjected to Illumina HiSeq sequencing (PE150).

### *In vitro* pre-crRNA processing assay

RNA processing assays were conducted in a 1×reaction buffer (50 mM NaCl, 10 mM Tris-HCl, 10 mM MgCl₂, and 1 mM DTT). 200 nM PmCas12m was complexed with 500 nM crRNA at 37 °C for 30 minutes. The reaction was incubated with Proteinase K (NEB) at 37 °C for 15 minutes, followed by adding 2x formamide stop buffer (90% formamide, 0.075% bromophenol blue, 0.075% xylene cyanol FF, and 50 mM EDTA) and running on a 15% TBE-urea-PAGE gel. Products were visualized by staining with SYBR Gold stain. The RNA templates used in this study are listed in Supplementary Table 5.

### *In vitro* DNA binding and cleavage assay

For the 6-FAM-labeled dsDNA binding assay, 100 nM PmCas12m was incubated with 250 nM sgRNA in 1× reaction buffer at 37 °C for 30 minutes. Subsequently, 20 nM substrate dsDNA was added, and the mixture was incubated for an additional 30 minutes at 37 °C. The reaction was quenched by adding 2× formamide stop buffer. The reaction products were resolved on a 6% DNA retardation gel at 120 V for 40 to 60 minutes. To determine whether PmCas12m lacks DNA cleavage activity *in vitro*, the ternary complex was formed as described above. The reaction was treated with RNase A (NEB) at 37°C for 15 minutes, followed by incubation with Proteinase K (NEB) at 37 °C for 15 minutes. The reaction was stopped by adding 2× formamide stop buffer, then heated to 95 °C for 5 minutes, and analyzed by electrophoresis on a 15% TBE-urea-PAGE gel. Sequence information for the substrates used in this study is listed in Supplementary Table 5.

### *In vivo* transcriptional silencing assay

The *in vivo* transcriptional silencing assay was performed as previously described with minor modifications.^13^ *E. coli* cells harboring the pCas12m and pCRISPR plasmids were co-transformed with pTarget plasmid. After recovery, the transformation mixture was inoculated into 3 mL of LB medium containing the appropriate antibiotics at 37°C for 16-20 hours. The cells were subsequently analyzed by FACS, qRT-PCR and transformation assays.

### Quantitative RT-PCR

Cells were harvested for total RNA isolation using the TransZol Up RNA Extraction Kit (TransGen) according to the manufacturer’s protocol. cDNA synthesis was performed using TransScript cDNA Synthesis SuperMix (TransGen). qRT-PCR reactions were prepared in 96 well plates with PerfectStart Green qPCR SuperMix (TransGen) and run on Archimed X4 system (RocGene). All the primers are listed in Supplementary Table 3.

### Lentivirus and AAV production

Lentivirus packaging was performed following the published protocol^61,62^. Briefly, HEK293T cells were seeded in 10 cm dishes and grown to approximately 80% confluency. Cells were co-transfected with 10 µg of the lentiviral DMS plasmid library, 2 µg of the VSV-G envelope plasmid, and 5 µg of the psPAX2 packaging plasmid using PEI transfection reagent. After 72 hours post-transfection, the culture supernatants containing lentiviral particles were collected, filtered through a 0.45 µm low protein-binding filter, and then stored at –80 °C for further use. The AAV-xCas12m-CRISPRoff-sgRNA vectors were packaged into AAV9 and produced and purified using a standard triple transfection as previously described^63^.

### DMS Library construction, FACS, and deep sequencing

By combining structural analysis, the DMS library of PmCas12m variants was divided into 4 units, and the saturation mutants were synthesized by Genscript based on the lenti-PmCas12m-VPR-RFP vector. To avoid swapping contamination, each of the four libraries was individually packaged into the lentivirus and subsequently transduced into TRE3G-GFP HEK293T cells that stably expressed the corresponding sgRNA. The MOI was less than 0.3 to ensure that no more than one mutant PmCas12m variant was expressed per cell. 72 hours post-transduction, the high GFP-expression cells and low GFP-expression cells were sorted and cultured for 5 days. After that, we re-sorted the high GFP-expression cells and low GFP-expression cells separately. The total number of collected cells was about 1 million cells per group. Total RNA was extracted from each group, followed by cDNA synthesis. Each library was subsequently amplified with primers containing specific adaptors and barcodes for Illumina HiSeq sequencing (PE150). The abundance of each mutation was counted and normalized to the total number of reads in each group, The activity was calculated as the ratio of high GFP-expression reads to low GFP-expression reads.

### Animal experimentation

All animal experiments were approved by the Institutional Animal Care and Use Committee of Peking University Health Science Center. Six weeks old C57BL/6 mice were injected with 1×10^10^ viral genome (vg) of AAV-HBV1.04 through the tail vein. After 7 days, these mice were randomly divided into three groups and injected with 2×10^11^ vg of AAV-xCas12m-CRISPRoff-sgNT, AAV-xCas12m-CRISPRoff-sg1 or AAV-xCas12m-CRISPRoff-sg9, respectively. Blood samples were collected at the indicated time points, and serum samples were obtained by centrifugation at 1500 g for 15 minutes at 4°C to measure the levels of HBV DNA, HBeAg, and HBsAg.

### Analysis of HBV translation and replication

The levels of HBeAg and HBsAg in cell supernatant and mouse serum were measured using an ELISA Kit (Autobio) according to the manufacturer’s protocol. HBV DNA quantification was performed as previously described with minor modifications^64^. In brief, HBV DNA was extracted from the cell pellets, supernatant or serum using TIANamp Virus DNA/RNA Kit (TIANGEN) and quantified by qRT-PCR. Viral genome equivalent copies were calculated using a standard curve generated with 2×HBV plasmid at concentrations ranging from 10^1^–10^9^ copies. For the detection of cccDNA, the extracted DNA samples were treated with Plasmid-safe DNase (Lucigen) at 37°C for 30min to degrade the linear DNA, followed by heat inactivation at 70°C for 30 minutes. The reaction products were subsequently quantified by qRT-PCR, with viral genome equivalent copies determined using a standard curve generated from known copy numbers. All primers used are listed in Supplementary Table 3.

---

**Extended Data Fig. 1.**
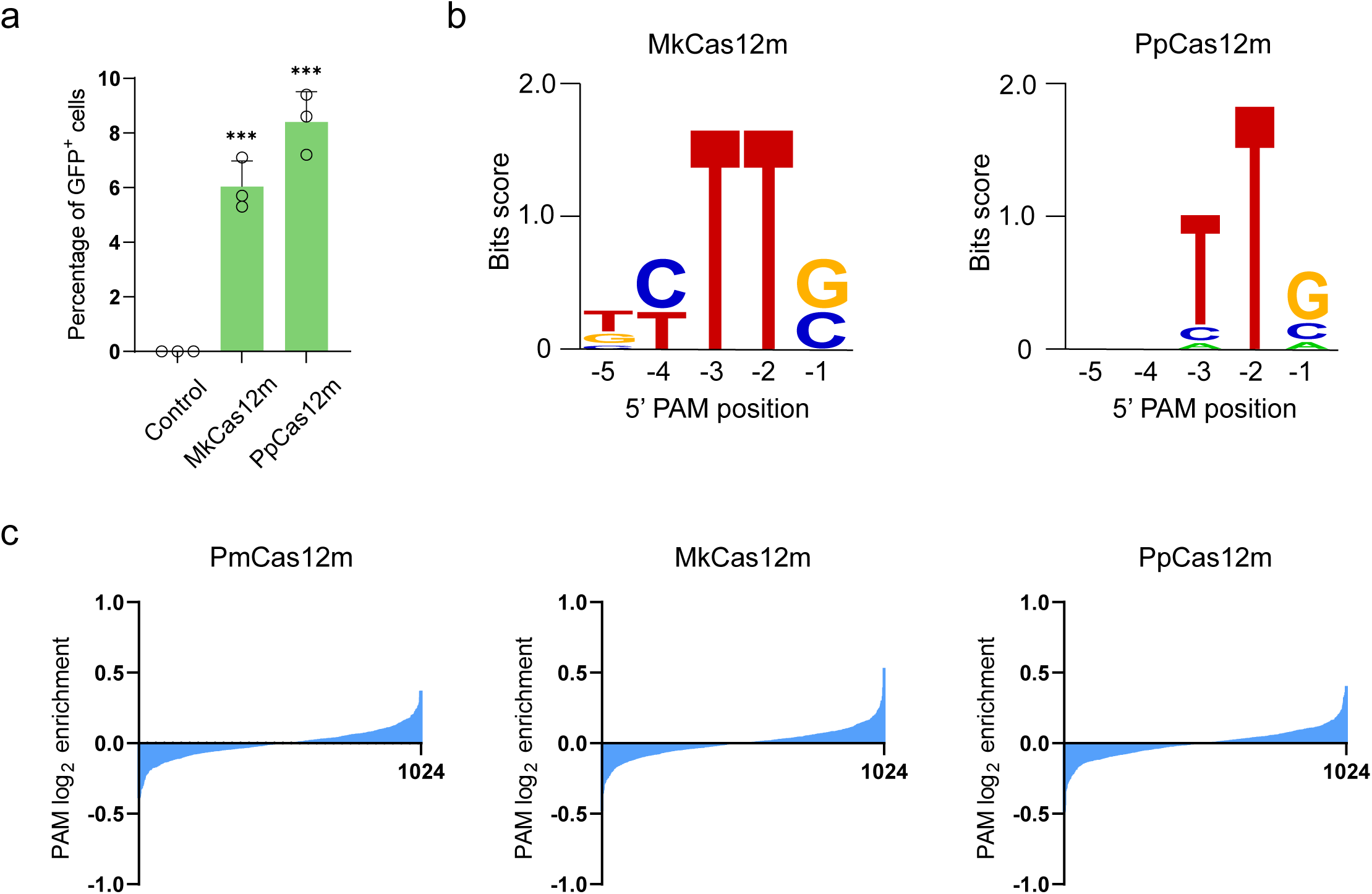
Identification of CRISPR-Cas12m homologs. **a**, FACS analysis of the GFP-positive cell percentage for MkCas12m and PpCas12m. Data represent the mean ± s.d. of three biological replicates. ***P value < 0.001. **b**, PAM motifs for MkCas12m (left) and PpCas12m (right). **c**, PAM depletion assay results for PmCas12m, MkCas12m and PpCas12m.

**Extended Data Fig. 2.**
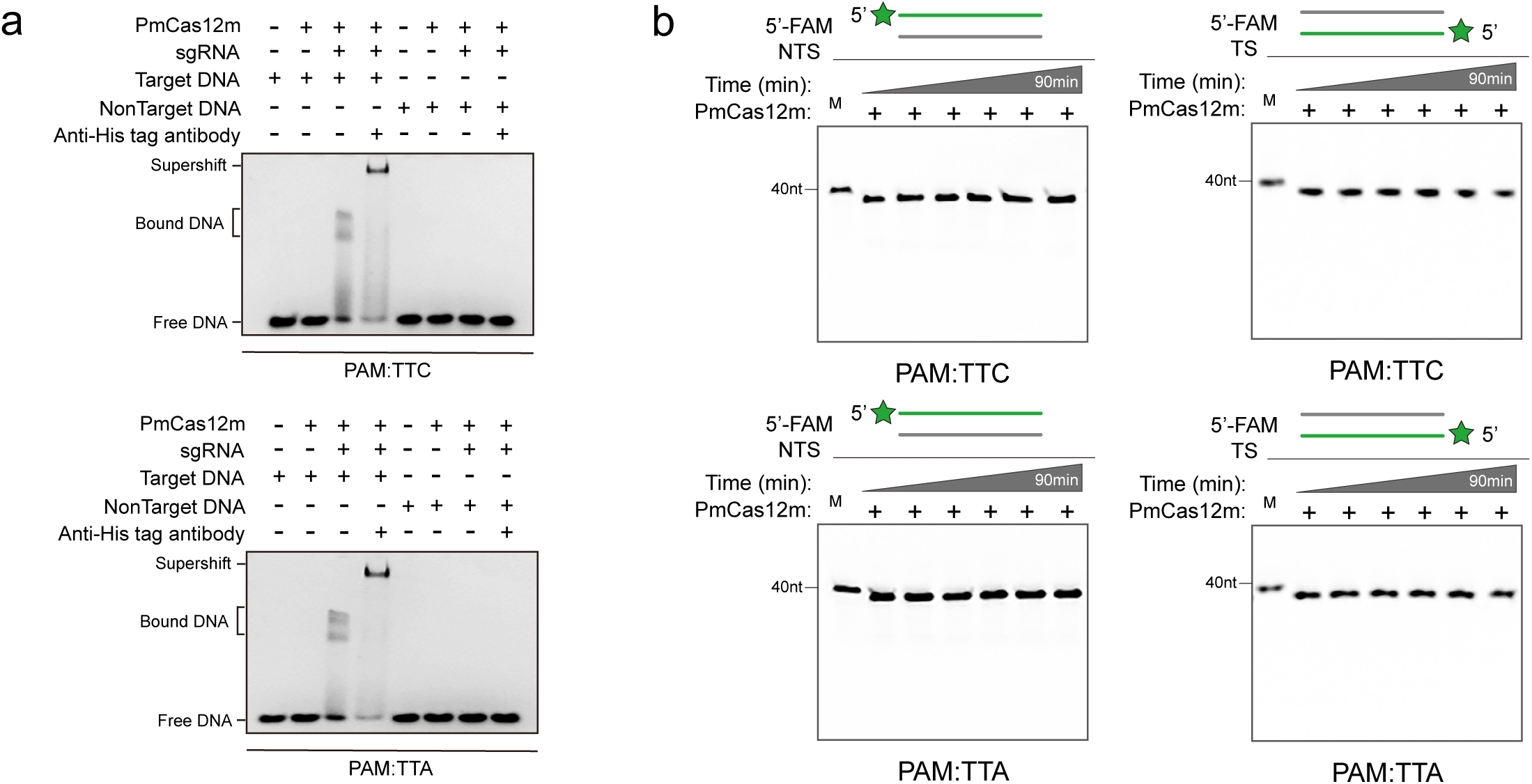
The nuclease-dead characteristic of PmCas12 *in vitro*. **a**, DNA binding assay demonstrates the DNA recognition capability of PmCas12m *in vitro*. **b**, DNA cleavage assay reveals the nuclease-deficient property of PmCas12m *in vitro*.

**Extended Data Fig. 3.**
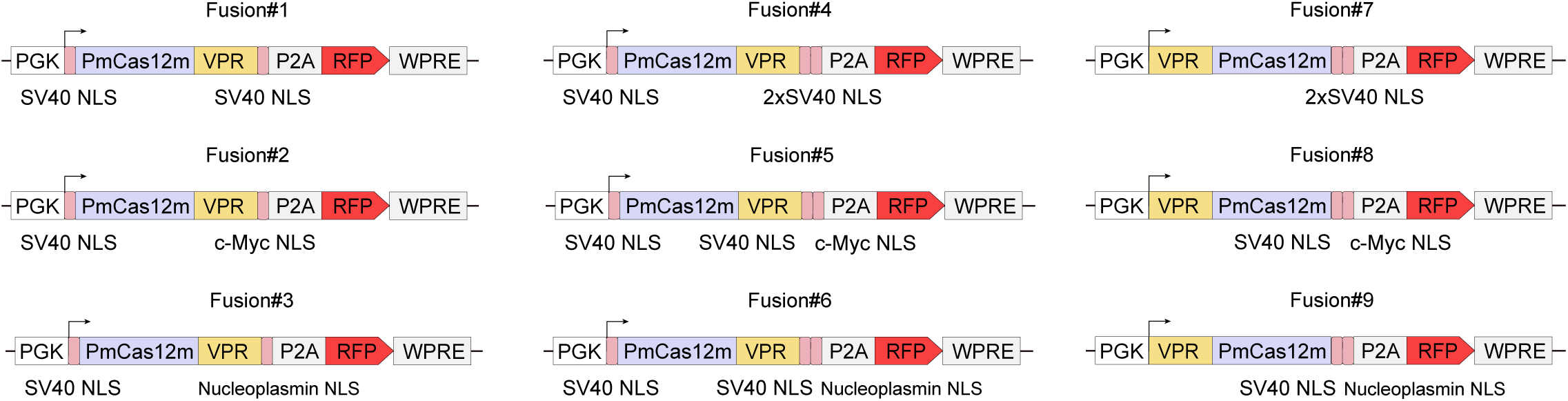
Schematic of PmCas12m-VPR fusion designs (#1-#9). The designs include PmCas12m fused to VPR at either the N-terminus or C-terminus, with nuclear localization signals (NLSs) from SV40, nucleoplasmin, or c-Myc.

**Extended Data Fig. 4.**
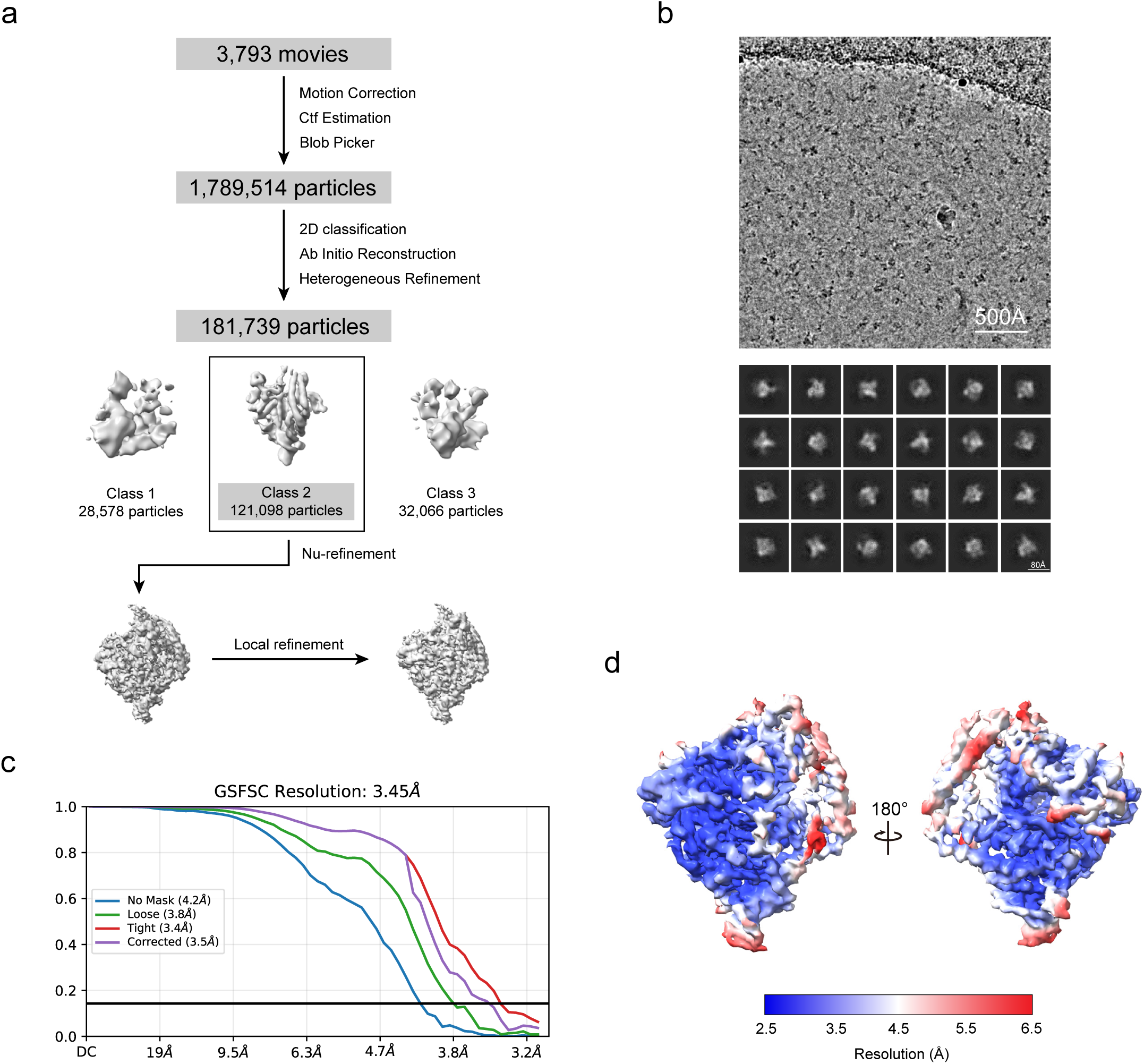
Cryo-EM data processing workflow for the PmCas12m-crRNA–target DNA ternary complex. **a**, Workflow for the single-particle cryo-EM image processing. **b**, Representative cryo-EM image and 2D class-averages of the PmCas12m-crRNA–target DNA complex. **c**, Euler angle distribution of refined particles used in generating the final map. **d**, Local resolution of the cryo-EM density map.

**Extended Data Fig. 5.**
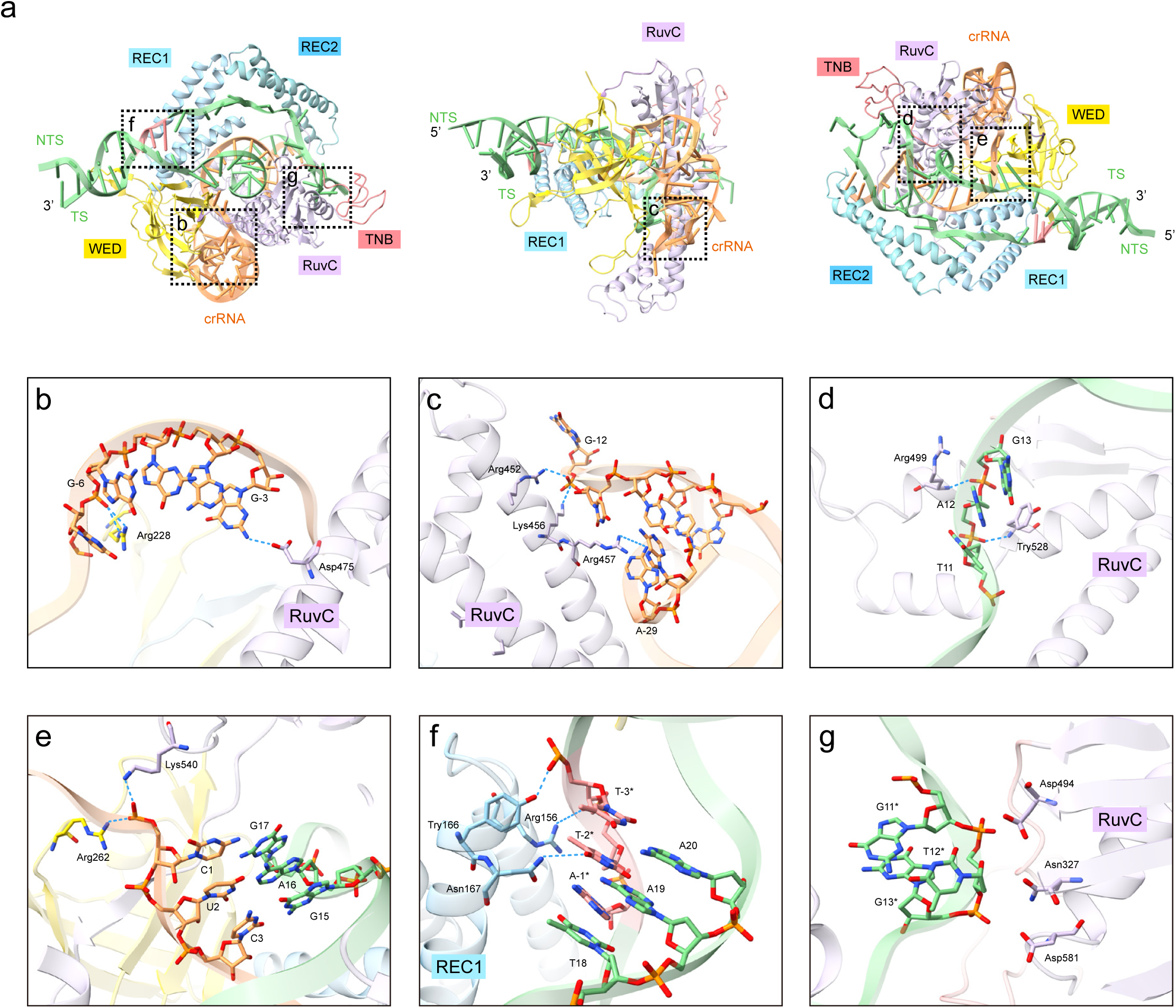
crRNA and target DNA recognition. **a**, Recognition sites of the crRNA and target DNA. **b-f**, Detailed recognition of crRNA **(b,c)**, target DNA**(d)**, DNA-RNA heteroduplex**(e)** and the PAM duplex**(f)**. **g**, Close-up view of the non-canonical RuvC residues.

**Extended Data Fig. 6.**
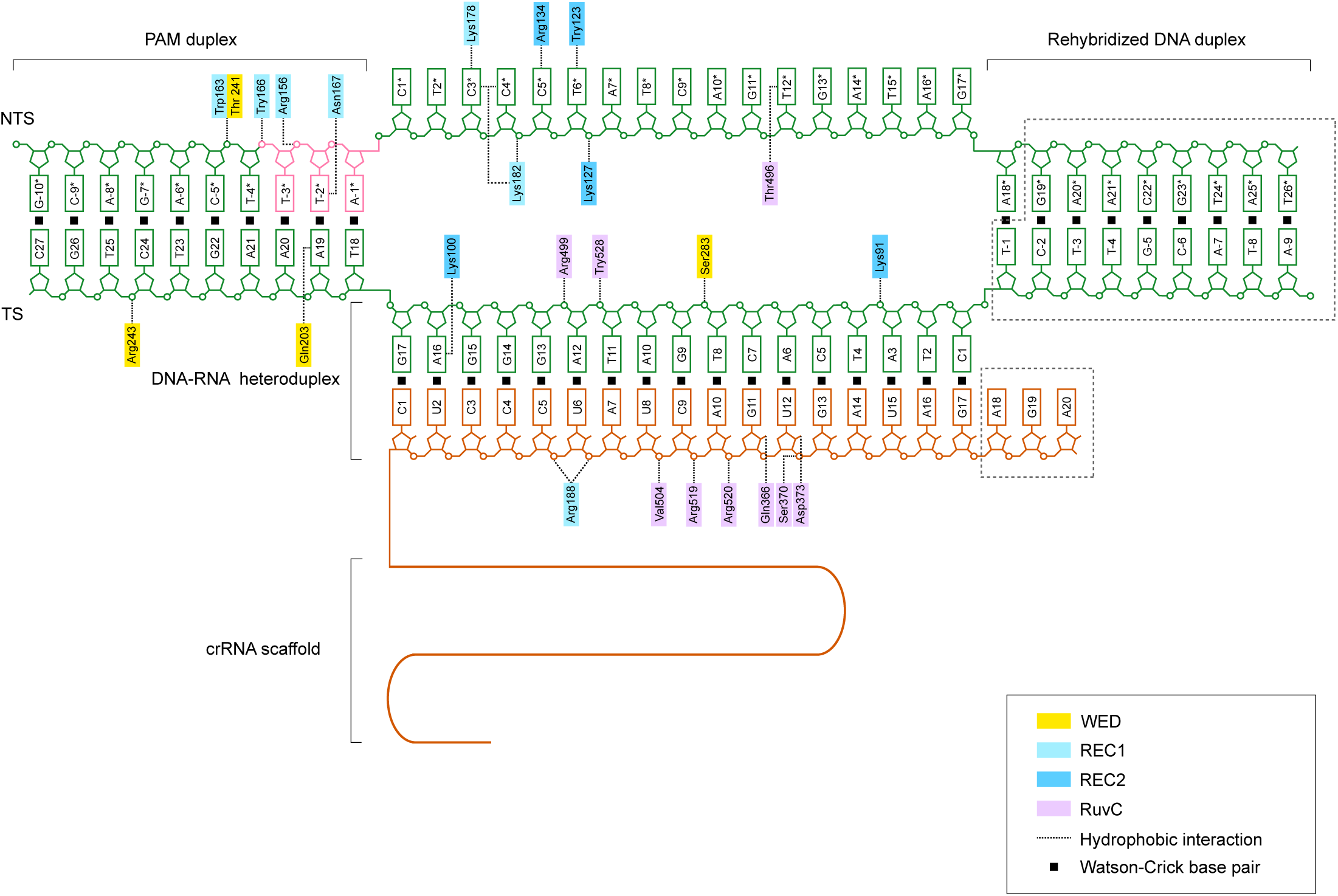
Schematic of nucleic acid recognition. Residues of PmCas12m that interact with nucleic acids are highlighted in their corresponding colors. Disordered regions are represented by dashed gray lines.

**Extended Data Fig. 7.**
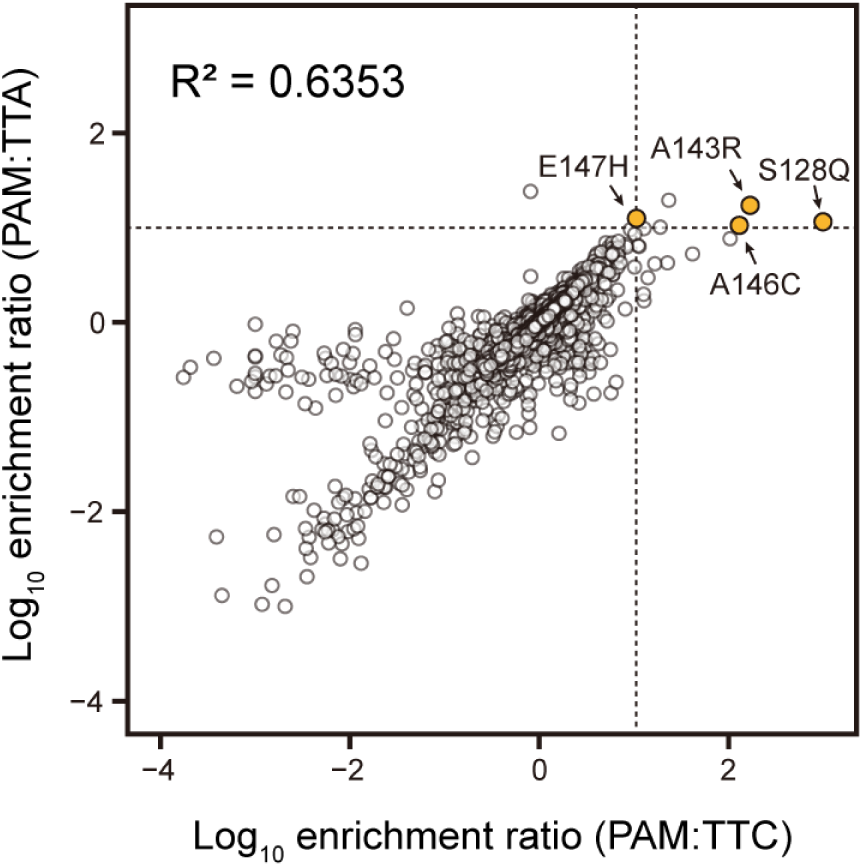
Correlation of epigenome editing efficiency using DMS method. Correlation of single-mutant effects on the epigenome editing efficiency of PmCas12m, with sgRNAs targeting the TRE3G promoter using TTA or TTC PAMs.

**Extended Data Fig. 8.**
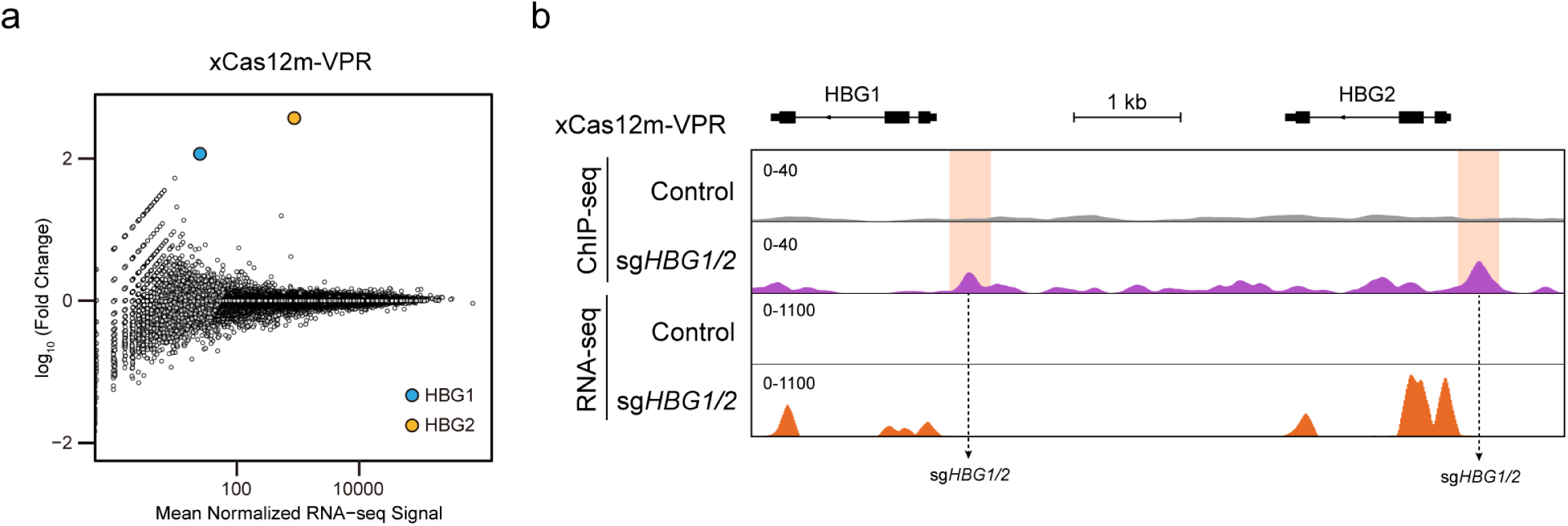
Characterization of xCas12m-VPR off-target effects in human cells. **a.**Volcano plots highlight the RNA-seq profiles of HEK293T cells transfected with xCas12m-VPR and sgRNA targeting the *HBG1/2* promoter, compared to cells transfected with non-targeting sgRNA. Data represent the mean of two biological replicates. **b**. Density maps of ChIP-seq (HA-tagged xCas12m-VPR) and RNA-seq at the *HBG1* and *HBG2* loci in HEK293T cells co-expressing xCas12m-VPR with sgRNA targeting the *HBG1*/*2* promoter, compared to cells transfected with non-targeting sgRNA.

**Extended Data Fig. 9.**
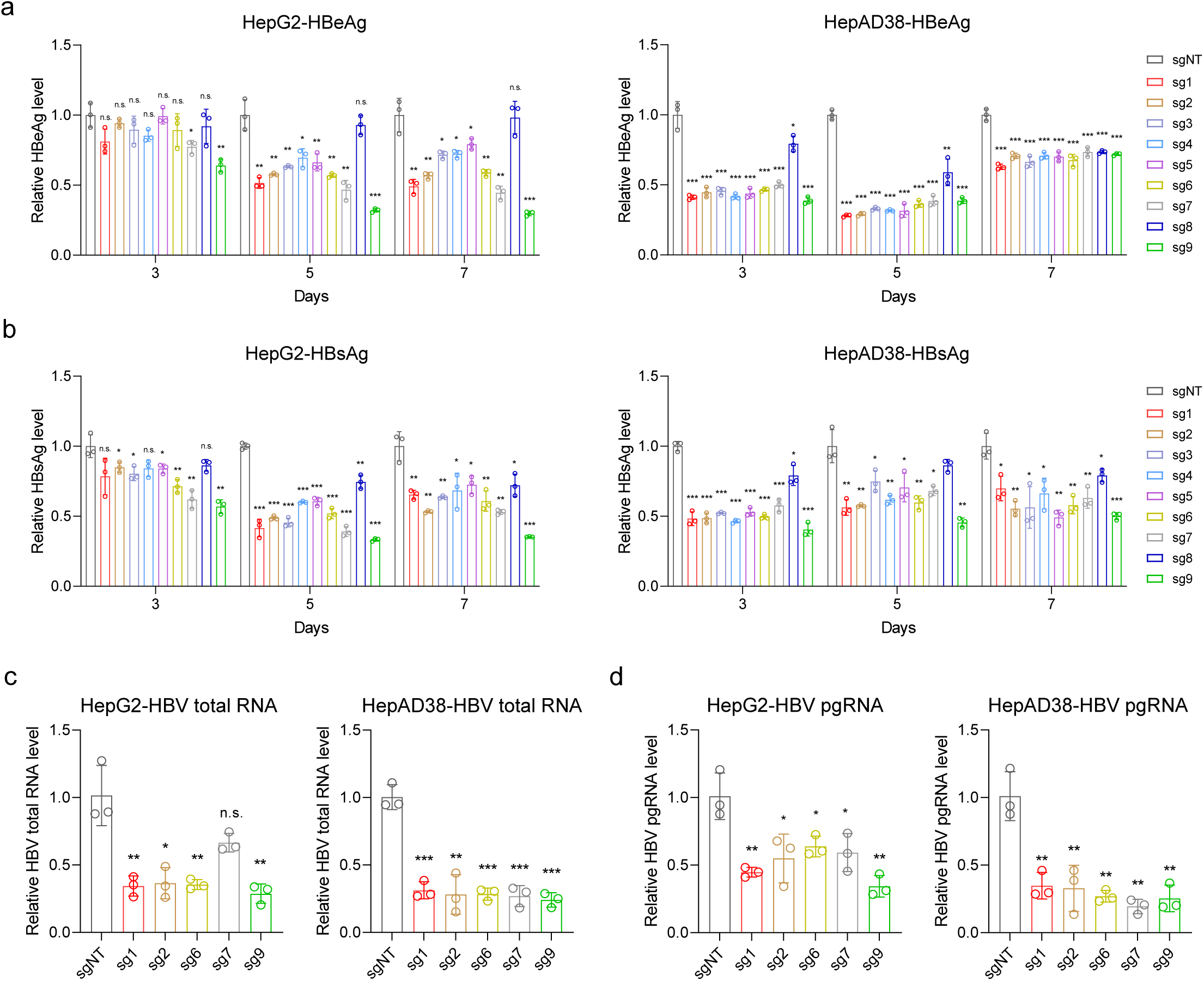
xCas12m-CRISPRoff-mediated epigenetic silencing suppresses viral product expression. **a, b** Levels of HBeAg(**a**) and HBsAg(**b**) in the cell culture supernatant of HepG2 cells (left) and HepAD38 cells (right) at the indicated time points post-transfection with xCas12m-CRISPRoff systems. Data are presented the mean ± s.d. of three biological replicates. ***P value < 0.001, **P value < 0.01, *P value < 0.05, n.s., non-significant. **c**, Total HBV RNA levels in HepG2 cells (left) and HepAD38 cells (right) on day 7 post-transfection with xCas12m-CRISPRoff systems. Data are presented the mean ± s.d. of three biological replicates. ***P value < 0.001, **P value < 0.01, *P value < 0.05, n.s., non-significant. **d**, Levels of pgRNA in HepG2 cells (left) and HepAD38 cells (right) on day 7 post-transfection with xCas12m-CRISPRoff systems. Data are presented the mean ± s.d. of three biological replicates. ***P value < 0.001, **P value < 0.01, *P value < 0.05, n.s., non-significant.

**Extended Data Fig. 10.**
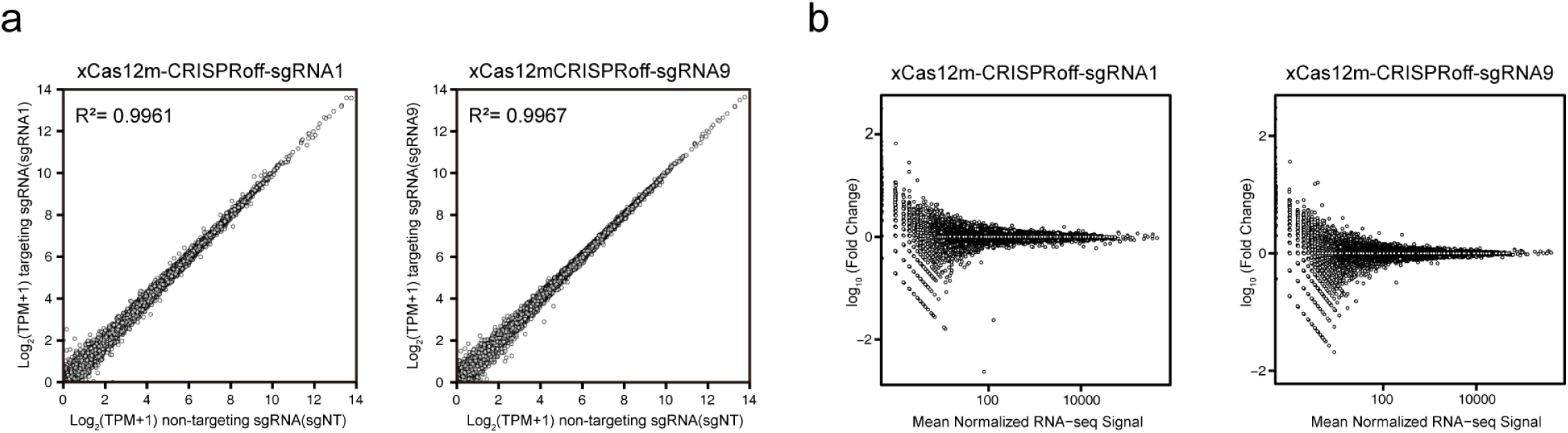
The impact of sgRNAs targeting the HBV genome on the human transcriptome in HBV-infected HepG2-NTCP cells. **a**, RNA-seq analysis of HBV-infected HepG2-NTCP cells targeting the HBV genome compared to non-targeting controls. TPM, transcripts per million mapped reads. Data represent the mean of two biological replicates. **b**, Volcano plots showing human transcriptome profiles with sgRNAs targeting the HBV genome. Data represent the mean of two biological replicates.

